# Why not record from *every* electrode with a CMOS scanning probe?

**DOI:** 10.1101/275818

**Authors:** George Dimitriadis, Joana P. Neto, Arno Aarts, Andrei Alexandru, Marco Ballini, Francesco Battaglia, Lorenza Calcaterra, Susu Chen, Francois David, Richárd Fiáth, João Frazão, Jesse P Geerts, Luc J. Gentet, Nick Van Helleputte, Tobias Holzhammer, Chris van Hoof, Domonkos Horváth, Gonçalo Lopes, Carolina M. Lopez, Eric Maris, Andre Marques-Smith, Gergely Márton, Bruce L. McNaughton, Domokos Meszéna, Srinjoy Mitra, Silke Musa, Hercules Neves, Joana Nogueira, Guy A. Orban, Frederick Pothof, Jan Putzeys, Bogdan C. Raducanu, Patrick Ruther, Tim Schroeder, Wolf Singer, Nicholas A. Steinmetz, Paul Tiesinga, Istvan Ulbert, Shiwei Wang, Marleen Welkenhuysen, Adam R. Kampff

**Affiliations:** Sainsbury Wellcome Centre for Neural Circuits and Behaviour, University College London, United Kingdom; ATLAS Neuroengineering, Leuven, Belgium; imec, 3001 Leuven, Belgium; Donders Institute for Brain, Cognition and Behavior, Radboud University Nijmegen, The Netherlands; Team Waking, Lyon Neuroscience Research Center (CNRL), INSERM-U1028, CNRS-UMR5292, Bron, France; Institute of Cognitive Neuroscience and Psychology, Research Centre for Natural Sciences, Budapest, Hungary; Faculty of Information Technology and Bionics, Pazmany Peter Catholic University, Budapest, Hungary; Champalimaud Neuroscience Programme, Champalimaud Centre for the Unknown, Portugal; School of PhD Studies, Semmelweis University, Budapest, Hungary; Institute for Integrated Micro and Nano Systems (IMNS), University of Edinburgh, UK; Uppsala University, Sweden; Department of Medicine and Surgery, University of Parma, Parma, Italy; Department of Microsystems Engineering (IMTEK), University of Freiburg, Germany; Electrical Engineering Department-ESAT, KU Leuven, 3001 Leuven, Belgium; Max Planck Institute for Brain Research, Frankfurt a. M., Germany; Neurinformatics department, Donders Institute for Brain, Cognition and Behavior, Radboud University Nijmegen, Heyendaalseweg 135, 6525 AJ, Nijmegen, The Netherlands; Fiocruz, Brazil; Departamento de Ciência dos Materiais, CENIMAT/I3N and CEMOP/Uninova, Faculdade de Ciências Tecnologia-Universidade Nova de Lisboa, Caparica, Portugal; Department of Neurobiology and Behavior, University of California at Irvine, Irvine CA, USA, 92697-8439; Department of Neuroscience, The University of Lethbridge, Lethbridge, AB, Canada, T1K 3M4; Cluster of Excellence BrainLinks-BrainTools, University of Freiburg, Germany; Department of Biological Structure, University of Washington, Seattle, WA; University College London; Janelia Research Campus, HHMI; Frankfurt Institute for Advanced Studies, Frankfurt a. M., Germany; Ernst Strüngmann Institute for Neuroscience in Cooperation with Max Planck Society, Frankfurt a. M., Germany

## Abstract

It is an uninformative truism to state that the brain operates at multiple spatial and temporal scales, each with each own set of emergent phenomena. More worthy of attention is the point that our current understanding of it cannot clearly indicate which of these phenomenological scales are the significant contributors to the brain’s function and primary output (i.e. behaviour). Apart from the sheer complexity of the problem, a major contributing factor to this state of affairs is the lack of instrumentation that can simultaneously address these multiple scales without causing function altering damages to the underlying tissue. One important facet of this problem is that standard neural recording devices normally require one output connection per electrode. This limits the number of electrodes that can fit along the thin shafts of implantable probes generating a limiting balance between density and spread. Sharing a single output connection between multiple electrodes relaxes this constraint and permits designs of ultra-high density probes.

Here we report the design and in-vivo validation of such a device, a complementary metal-oxide-semiconductor (CMOS) scanning probe with 1344 electrodes; the outcome of the European research project NeuroSeeker. We show that this design targets both local and global spatial scales by allowing the simultaneous recording of more than 1000 neurons spanning 7 functional regions with a single shaft. The neurons show similar recording longevity and signal to noise ratio to passive probes of comparable size and no adverse effects in awake or anesthetized animals. Addressing the data management of this device we also present novel visualization and monitoring methods. Using the probe with freely moving animals we show how accessing a number of cortical and subcortical brain regions offers a novel perspective on how the brain operates around salient behavioural events. Finally, we compare this probe with lower density, non CMOS designs (which have to adhere to the one electrode per output line rule). We show that an increase in density results in capturing neural firing patterns, undetectable by lower density devices, which correlate to self-similar structures inherent in complex naturalistic behaviour.

To help design electrode configurations for future, even higher density, CMOS probes, recordings from many different brain regions were obtained with an ultra-dense passive probe.

## Introduction

The number of neurons that can be monitored simultaneously during an extracellular recording is currently limited by the number of electrodes that can be implanted (Stevenson, 2017; Stevenson and Kording, 2011). However, the desire for large-scale monitoring of single neuron activity and the need to minimize tissue damage compete with one another (Buzsáki et al., 2015). The high resolution processes of semiconductor fabrication (i.e., photolithographic patterning of thin film conductors and insulators on a silicon substrate) have enabled dozens of microelectrodes, packed with an ever increasing density, along needle-like probes (Blanche et al., 2005; Dugué et al., 2011; Torfs et al., 2011; Berenyi et al., 2014; Buzsáki et al., 2015; Shobe et al., 2015; Herbawi et al., 2018; Dorigo et al., 2018). In the standard “passive” silicon probe, electrodes distributed along the probe shaft must each be connected to an external contact pad on the probe base by metal lines deposited along the shaft. Advances in lithography techniques (i.e., e-beam lithography (Rios et al., 2016; Scholten and Meng, 2016)), metal deposition procedures, and quality control have made it possible to fabricate a five shank, 1000-channel probe, where each shank (width of ~50 μm) has 200 electrodes of 9 × 9 μm and a pitch of 11 μm (Scholvin et al., 2016). Despite these impressive designs, a bottleneck remains in the number of output connection lines that can be squeezed into a single, thin probe shaft. Therefore, monitoring the thousands of electrodes required to densely cover an 8 mm × 100 μm probe will demand a new technology; a technology that multiplexes the signal from multiple electrodes into one output line.

The semiconductor industry has solved the same interconnect challenge with integrated circuitry (IC), often fabricated using the CMOS manufacturing process. As a characteristic example, mobile phones with on-board cameras use CMOS image sensors (CIS) that are lightweight and low power, but that can convert an optical image into an electronic signal with very high spatial resolution. A CIS is composed of an array of millions of identical pixels, each having at least a photodiode, an addressing transistor that acts as a switch, and an amplifier. Each pixel is sampled once within each frame and the image output is generated by rapidly scanning the full sensor matrix. This active switching / time multiplexing is crucial for achieving the small size of a CIS device, as it conveys the signals from many pixels with just a few output connections.

In the case of silicon probes, their one-dimensional architecture, high sampling speed requirements (three orders of magnitude larger than a CIS), the low amplitude of the neural signals (three orders of magnitude lower than the photodiode signals in CMOS cameras) and the need to avoid tissue overheating due to electrical power dissipation have challenged the inclusion of integrated circuitry within the probe’s shaft. The concept of integrating electronic elements in the same substrate used to build the recording electrodes was first introduced in the 1980s (Najafi and Wise, 1986; Najafi et al., 1985; Wise and Ji, 1992; Wise and Najafi, 1991) where microelectronic components have been integrated in the probe base. This was followed by the European project *NeuroProbes,* pioneering the CMOS integration into the slender probe shaft introducing the electronic depth control (EDC) approach (Neves et al., 2008a; Seidl et al., 2011; Torfs et al., 2011). The most recent CMOS-based implantable probe designs for *in vivo* applications integrate circuitry into both the probe base and within the probe shaft(s) and have been reviewed elsewhere (Lopez et al., 2013, 2016; Obien et al., 2015; Ruther and Paul, 2015; Raducanu et al., 2016; Seymour et al., 2017; Steinmetz et al., 2018).

The NeuroSeeker European research project aimed to produce CMOS-based, time multiplexing, silicon probes for extracellular electrophysiological recordings with 1344 electrodes on a 50 μm thick, 100 μm wide, and 8 mm long shank with a 20 × 20 μm electrode size and a pitch of 22.5 μm (see Box 1 for a technical overview of the probe and (Raducanu et al., 2016, 2017) for an in depth description of the probe’s design). This innovative solution has allowed the neuroscientists within the consortium to record brain activity with an unprecedented combination of resolution and scale. Here we report the use of the NeuroSeeker probe to record from both anesthetized and freely moving rats. We characterize in saline and *in vivo* the increase in noise that the local amplification and time multiplexing scheme incurs relative to a passive electrode probe with the same configuration. We show that this increase is insignificant *in vivo* and does not inhibit the capture of high quality neural signals. In the case of freely moving animals, we also include data showing that the signal remains stable for at least 10 days. We also describe a method for removing the NeuroSeeker probe intact, and ready for reuse, from the brain of a chronically implanted animal and show results from freely moving recordings from two animals implanted with such a ‘second hand’ probe. This allows the multiple re-use of probes whose price, and low production numbers, makes them currently inaccessible to all but a tiny minority of electrophysiology labs around the world.

Recording with this number of electrodes results in datasets of around 200 GB per hour with spike numbers in the millions. This scale of data poses new difficulties in the visualization, pre-processing, and curation of the signal, even before it can be used to explore correlations with sensory stimuli or behaviour. Here we present an online visualization method that allows for the rapid overview of the signal from all 1344 electrodes, giving the experimenter a global picture of the recording quality as data is acquired.

Using a CMOS probe with very high count and density of recording channels allows accessing an unprecedented number of neurons from a large and distributed number of cortical and subcortical brain regions. It is unclear though how such information can be analysed and, given its complexity, actually lead to insights in the way the brain encodes its environment and produces behaviour. Results of recordings from such scales have started to alter the way the community thinks about brain regions and inter-regional communication (Petreanu et al., 2012; Stringer et al., 2019; Wekselblatt et al., 2016). Most of these paradigm shifting efforts though record only from cortical regions or from a limited number of subcortical ones due to the limitations of the recording toolboxes used up to today. We show how the NeuroSeeker probe’s design adds a tool to these toolboxes that allows recordings from multiple regions, with high counts of neurons from freely moving animals and with minimal damage to brain tissue (one vs. multiple shafts). More specifically we demonstrate that probes that span multiple areas can capture brain behaviour around salient moments of an animal’s behaviour that would be missed by probes with smaller brain coverage. We then compare the NeuroSeeker signal with signals of equal spread but sparser electrode distributions. This showcases how combining an increase in electrode density and in channel count leads to detection of neuronal firing patterns that much better correlate to behavioural patterns found in complex, freely moving, naturalistic behaviour.

Given that CMOS probes with local amplification and time multiplexing allow a significant increase in electrode density without sacrificing compactness, the neuroscientific community must now determine the physiologically relevant limit for the size and density of the electrodes on such probes. To help address this question, we have collected datasets from ultra-high density (256 electrodes of 5 × 5 μm size with a 6 μm pitch) passive probes. These were recorded from anesthetized rats in a wide range of brain structures and probe orientations. These datasets are available online for further analysis of the optimal geometries of future CMOS probes.

### Box 1

NeuroSeeker CMOS-based probe.

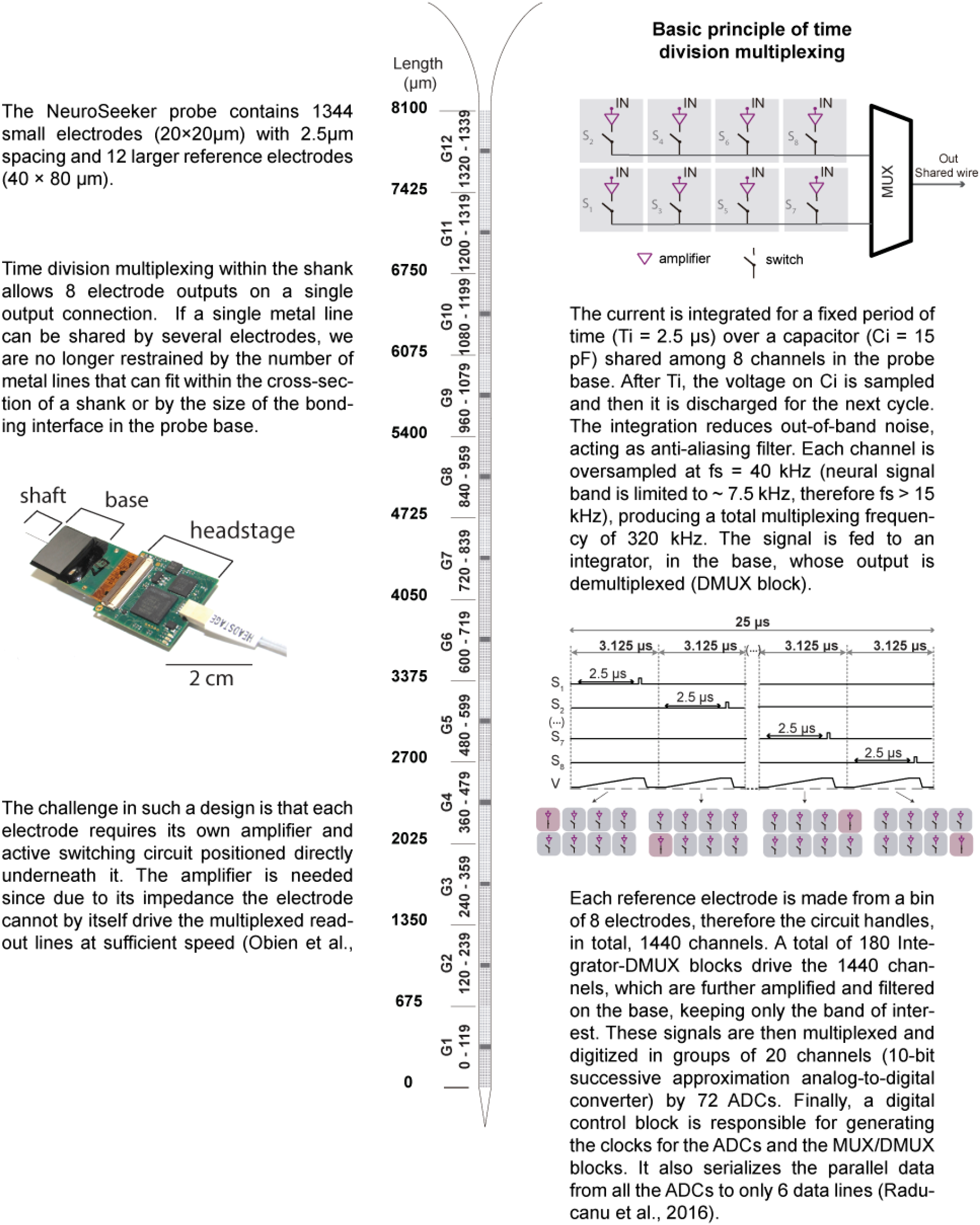

## Results

Several recordings were performed with the NeuroSeeker CMOS-based scanning probes *in vivo* to evaluate the viability of these devices. There are 12 groups of electrodes on the probe, which can be individually powered, labelled from G1 to G12, in Box 1 and Figure 1. Each group contains 120 channels: 112 electrodes and 1 local reference in the middle made from a group of 8 electrodes binned together. For each electrode, the user can select the reference type (external or internal), gain, and frequency bandwidth (high-pass cut-off frequency in action potential (AP) mode or low-pass cut-off frequency in local field potential (LFP) mode). For most recordings (unless otherwise noted), 1277 electrodes were set in the AP band (0.5–7.5 kHz), while 67 were set in the LFP band (1–500 Hz). The LFP electrodes were spaced 20 electrodes apart from one another, which translates to 5 rows, or, given the 22.5 μm pitch of each electrode, 112.5 μm.

**Figure 1.**
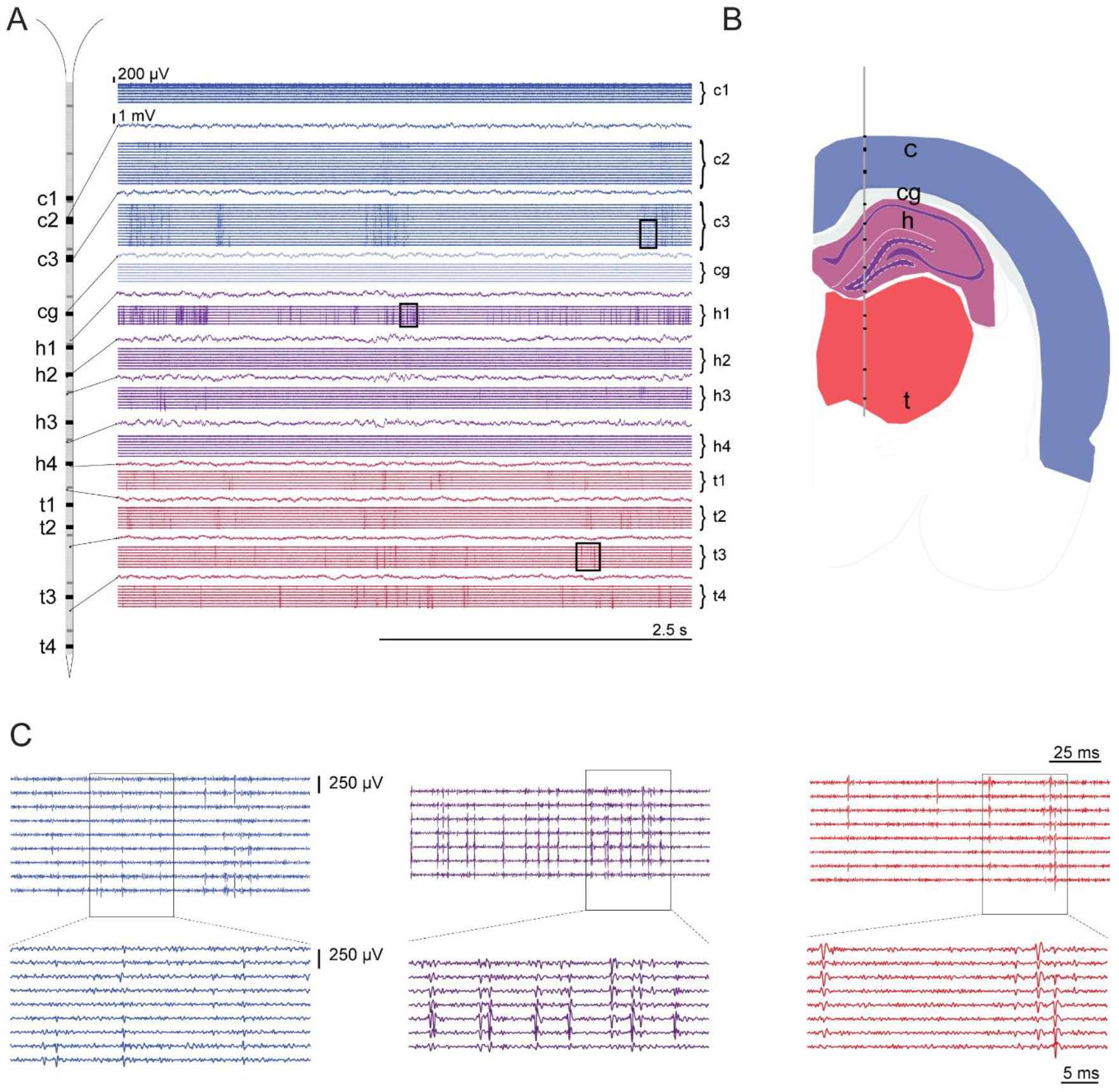
Example of a recording performed with 1120 electrodes simultaneously within cortex, hippocampus, and thalamus from an anesthetized rat. A) 5-s-long LFP and AP traces of a probe spanning multiple brain regions; B) Schematic of coronal slice signalling the group’s position shown in A as black blocks. ‘c, ‘cg’, ‘h’ and ‘t’ denote the anatomic locations of cortex, cingulum, hippocampus, and thalamus, respectively. The estimated probe position was based on the insertion coordinates as described in the Animal recordings part of the Methods section. From the point of insertion to the tip, the recording depth was 6.7 mm; C) Short epochs of AP band traces from groups c3, h1, and t3 (color-coded), illustrating the presence of diverse spike waveforms on several electrodes. These sections are highlighted in A by boxed areas on top of traces.

### Neural recordings

#### Acute recordings

Figure 1A shows a representative epoch of a recording performed simultaneously within cortex, hippocampus, and thalamus from an anesthetized rat. In this recording, 10 of the 12 groups are in the brain and thus enabled (powered on): 1060 AP mode and 60 LFP mode ones. A subset of traces from electrodes set in AP and LFP mode are shown in Figure 1A (a recording segment of all traces is shown in Supplementary Figure 1). The traces in AP mode are displayed in groups (c1, c2, c3, cg, h1, h2, h3, h4, t1, t2, t3 and t4) that correspond to the location on the probe shaft indicated by the black region in the probe schematic (Figure 1B). Moreover, traces of 11 out of 60 LFP mode electrodes are plotted between the group traces. The presence of fast voltage deflections in several traces reveals that the spiking activity of multiple neurons has been detected.

To evaluate if the data quality deteriorates with increasing number of groups powered, we computed the root mean square (RMS) noise of 212 electrodes (from groups G1 and G2 as defined in Box 1) set in AP mode while enabling more groups along the shank. The noise level for the 212 electrodes using the external reference (a screw in the animal’s skull), with 2 groups and 12 groups powered, is 10.2 ± 0.1 μV and 12.0 ± 0.1 μV, respectively. This compares to the noise reported by (Raducanu et al., 2017) of 12.4 ± 0.9 μV for 6 groups on. In that work the noise for a 12 group recording is not reported but data are shown that describe how a full probe recording (12 groups) has a low enough noise to allow recordings from hundreds of neurons (results that we also corroborate here). A point to consider is that the NeuroSeeker probe allows the calibration by the experimenter of the voltage supplied to the active electrodes. Lowering the number of on channels requires a smaller voltage which leads to a decrease in power dissipation and thus to a lower temperature increase along the probe’s shaft and base. In the Raducanu at al. results, the voltage supplied was kept to a value that would lead to an increase of the probe’s surrounding tissue temperature of no more than 1°C. In our work we did not adhere to this limitation, judging the quality of our recordings solely by the number of units we could detect. Further research should be done to elucidate how power dissipation affects the surrounding tissue, especially with tools that offer the experimenter the ability to balance number of electrodes with signal to noise and power dissipation.

The noise magnitude was also computed for the 1012 channels in the brain (half of group 10’s electrodes are outside of the brain) using either the external or the internal reference configuration while recording. The measured noise using the external (skull screw) and internal reference is 11.7 ± 0. 1 μV and 12.5 ± 0.1 μV, respectively. The noise magnitude computed across these datasets revealed no significant variability. We also measured the noise in saline for the NeuroSeeker probe. The measured noise in saline while powering the same 1012 channels was 9.4 ± 0.1 μV. We compared the noise measured in saline and during acute recordings with a 128-channel probe previously described (Neto et al., 2016). This passive electrode probe has the same configuration, electrode size, and electrode material (titanium nitride) as the CMOS-based probe (impedance magnitude at 1 kHz is around 50 kΩ). The noise value calculated with the 128-channel probe in saline, on data sets filtered with a high pass of 500 Hz, was 2.69 ± 0.02 μV and *in vivo (amplifier2015-08-28T20_15_45* in http://www.kampff-lab.org/validating-electrodes/) was 10.40 ± 0.04 μV. Despite the difference in saline (9.4 vs 2.7 μV) the noise difference *in vivo* was small (11.7 vs 10.4 μV) between the CMOS-based probe and the passive 128-channel probe, highlighting the dominance of “biological” noise for *in vivo* extracellular recordings.

Several recordings were performed with different reference configurations and number of active groups. These datasets are available online (http://www.kampff-lab.org/cmos-scanning/) and summarized in Supplementary Table 1. Figure 1 is derived from one of these recordings, *18_26_30.bin.*

#### Chronic recordings

We also conducted a set of recordings from chronically implanted, freely moving animals. Given the high cost and low availability of high density probes, we designed a method that allowed us to extract the CMOS probe from the brain of a chronically implanted animal at the end of the experimental period so that it can be reused for multiple experiments. This required a specially designed probe holder and a series of steps after animal euthanasia to ensure the probe is removed without breaking the fragile, 100-μm-wide shaft. The holder is a two-part design and is shown in Figure 2A, while in Methods we describe a detailed probe recovery technique. We have managed to successfully remove four out of five chronically implanted probes (two dummy and two fully functional). In the case of the unsuccessful attempt (up to that point, a fully functional probe), the probe holder exhibited a small angle deflection off the Dorsal-Ventral / Anterior-Posterior plane, even before cutting of the resin, indicating that the probe had already moved and thus broken while the animal was alive.

**Figure 2.**
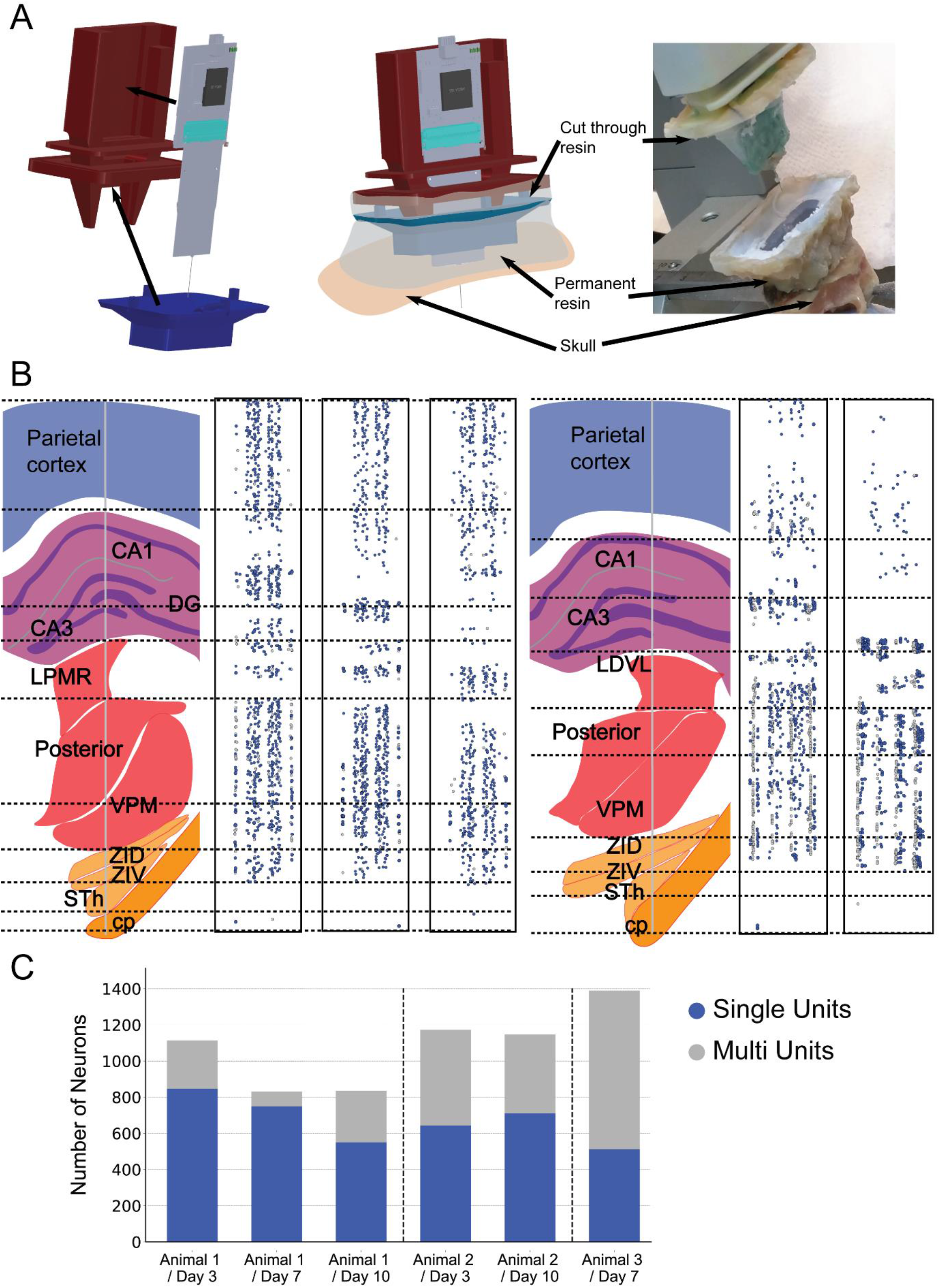
Chronic recordings and probe removal process. A) The processes of inserting the probe before surgery (left), implanting and gluing the probe in place during surgery (middle) and removing the probe by cutting the cut-through resin layer after surgery (right). B) Results from two animals chronically implanted with the same probe (Animals 1 and 2 implanted in that order). The anatomy schematics show where the probe was implanted. The scatter plots show the position of the recorded units on the probe, each for one recording session. Each dot is a single (blue) or a multi unit (grey) group and the axis are the y and x coordinates of the probe’s shaft (not shown to the same scale due to the extremely elongated nature of the shaft). The three position plots of the first animal show recordings done 3, 7 and 10 days after surgery, while the plots for the second animal show the day 3 and day 10 post surgery recordings. C) Cumulative bar chart showing the number of recorded units over days for three animals (Animals 1 and 2 are the same in B and C while Animal 3 was implanted after Animal 2 with the same probe). Brain regions: CA1: Cornu Ammonis 1, CA3: Cornu Ammonis 3, DG: Dentate Gyrus, LPMR: Lateral Posterior MedioRostral thalamic nucleus, LDVL: Lateral Dorsal VentroLateral thalamic nucleus: VPM: Ventral Posterior Medial thalamic nucleus, ZID: Zona Incerta Dorsal, ZIV: Zona Incerta Ventral, Sth: Subthalamic nucleus, cp: cerebral penducle

We have used freely moving recordings from three animals, all implanted with the same probe, to describe the long term recording capabilities and robustness in removal and re-implantation of the probe. In all cases the probe was fully implanted in the brain targeting the parietal cortex and the subcortical regions beneath (see Figure 2B schematics for a description of the anatomical regions the probe was passing through). For the first two animals this resulted in recordings from 1277 electrodes from the AP band and 67 electrodes from the LFP band. In the third animal we switched off the bottom two regions of the probe (which were showing too large a signal to noise ratio to be useful) thus creating recordings from 1064 AP electrodes and 56 LFP ones. In all cases, the probe was left in the animal for 4 weeks. Each animal was recorded daily in a freely moving session which ranged between 40 to 70 minutes. Each session generated between 3.3M and 8M spikes per hour of recording. Spikes were represented on groups of adjacent electrodes that ranged in size between 10 and 50 with most of the spikes showing on at least 40 electrodes (for a more detailed description of a neuron’s spread over the probe see (Raducanu et al., 2017)). We spike sorted some representative sessions to demonstrate the robustness of the recordings over time and our ability to remove the probe from one animal, implant it in another, and still get recordings of the same quality as a new probe, evidenced by the number of single units we could recover.

During the spike sorting pipeline (using Kilosort (Pachitariu et al., 2016) and t-SNE (Dimitriadis et al., 2018)) we separated the units into two categories. These were the single units and the multi units, i. e. the groups with spikes that originated from more than one neuron. Figure 2B shows the position of these units (color coded according to their assignment) on the probe shaft (see Methods on how these positions are calculated). For the first implanted rat (Animal 1) we show three recordings, taken 3, 7 and 10 days after surgery. For the second rat (Animal 2) we show days 3 and 10 post surgery (the choice of the recording days analysed was partially influenced by the quality of the animals’ behavioural output which allowed us to use the same data sets for multiple analysis). Figure 2C shows the number of the different types of units over the different recordings also showing results from a third rat (Animal 3, 7 days post surgery). The three animals were implanted sequentially with the same probe. Table 1 summarizes the number of units and number of spikes recorded in each session. The number of detected templates (most of them single units) ranged between 1389 and 831. In (Dimitriadis et al., 2018) we reported that in an under anesthesia session (with a different NeuroSeeker probe) and with 908 electrodes in the brain we detected 579 units; a number comparable to the unit count we get in these awake experiments. The data show that the probe proved to be a robust recording device, able to detect over 1000 units simultaneously while not showing a worse deterioration of its capacity compared to any other silicon probe over extended periods of recording.

**Table 1.** A summary of the number of units and number of spikes recorded during six representative sessions

|  | Animal 1/<br>Day 3 | Animal 1/<br>Day 7 | Animal 1/<br>Day 10 | Animal 2/<br>Day 3 | Animal 2/<br>Day 10 | 3<sup>rd</sup> Animal /<br>Day 7 |
| --- | --- | --- | --- | --- | --- | --- |
| Single units | 846 | 750 | 528 | 643 | 711 | 513 |
| Multi units | 267 | 81 | 320 | 529 | 436 | 876 |
| Total units | 1113 | 831 | 848 | 1172 | 1147 | 1389 |
| Spikes | 3,586,965 | 2,610,528 | 3,426,401 | 5,814,970 | 9,684,804 | 22,155,583 |
| Recording time | 40:30 | 46:58 | 56:53 | 46:20 | 1:14:34 | 2:36:24 |
| Spikes / hour | 5,380,447 | 3,332,588 | 3,636,596 | 7,543,744 | 7,816,629 | 8,521,378 |

### Data visualization

The signal traces presented in Figure 1 are just a small fraction of the total number of electrodes (105 of 1060 electrodes) recorded during an acute experiment (see all voltage traces in Supplementary Figure 1). These ultra-high channel count CMOS scanning probes presented a new challenge for the visualization and analysis of the acquired data. This challenge first appears when the user wants to monitor an ongoing recording, either to assess the signal quality appearing on each electrode or to better position the probe within the desired brain structure(s). Visualizing 1344 time series is difficult, even on an high definition (HD) monitor with 1080 rows of pixels. Furthermore, rendering voltage vs. time traces, the conventional representation for online physiology signals, is computationally expensive. However, modern graphic processors (GPUs) were specifically designed to parallelize such visualization tasks. We therefore developed a custom visualization pipeline using Bonsai (Lopes et al., 2015) by transferring each buffer of acquired probe data directly to the GPU and then using fragment shaders to render all probe data in real-time. The visualization shown in Figure 3 is a screen capture of one such online visualization in which the voltage signal of each channel is rendered from red to blue (also see Supplementary Movie 1). A neuron’s extracellular action potential is detected on many adjacent electrodes, and thus spikes in this visualization appear as a coloured block of red and blue stripes.

**Figure 3.**
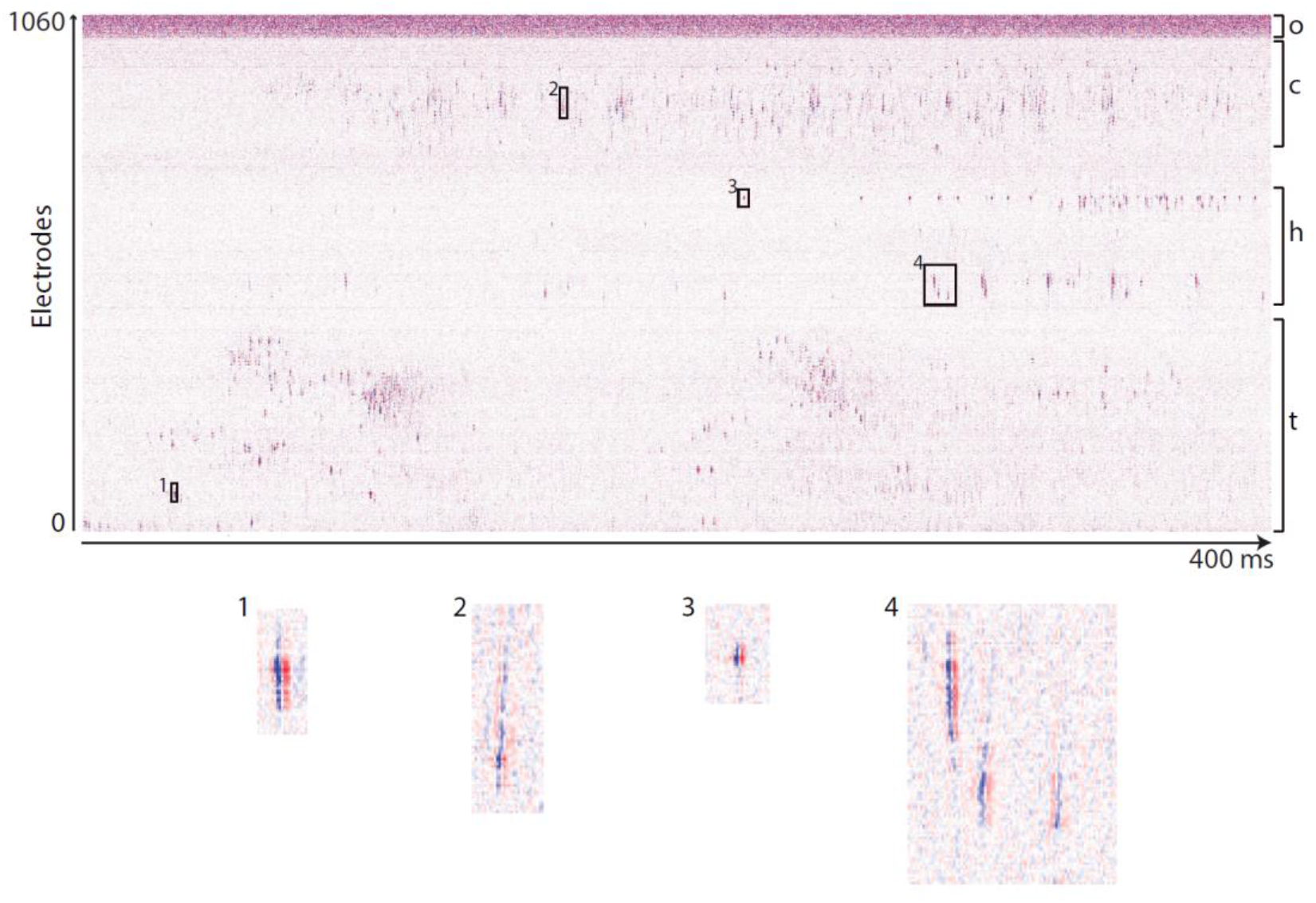
Real time GPU-based visualization of the voltage traces from 1060 electrodes in AP mode during acute recordings under anesthesia. The voltage traces recorded by each electrode (row) are displayed as a color saturation for each 100 microsecond bin (column). Red represents positive voltages (maximum color value represents 150 μV), blue negative voltages (minimum color value represents −150 μV), and white represents voltages near zero. This image shows 400 milliseconds of data recorded simultaneously in cortex, hippocampus and thalamus. ‘o’: out of the brain, ‘c’: cortex, ‘h’: hippocampus and ‘t’: thalamus.

### Global vs. local signals

Given a functional, high density, multi region covering probe we addressed the original NeuroSeeker consortium’s hypothesis that the signals recorded from this device in freely moving animals would offer a unique view of the brain’s activity combining local and global scales of activity.

The animals in our chronic recordings were engaged in a foraging task which at the end of each successful trial rewarded them with a chocolate pellet they could access from a nose poke. During their free foraging (in a 1 × 1 m arena) they were allowed to poke whether they had just finished a successful trial or not (generating both reward and exploratory pokes). At the same time our setup recorded them using a 120 fps camera. The animals were not deprived, thus generated a complex set of behaviours not overpowered by an overwhelming drive for reward.

### Global view of salient events

We used this setup to answer whether having access to both a large number of brain regions and to the detailed single neuron spiking information would be advantageous to our understanding of how the brain generates behaviour. We first focused on brain activity around the time of the pokes (in this case, the exploratory ones which were more numerous than the rewarded ones) and asked if the multi-regional coverage of the NeuroSeeker probe offered any advantage. The results for all animals can be seen in Figure 4. The figure shows the average over pokes (83, 96 and 100 pokes respectively) firing rates of all neurons, each normalised to itself, over a period of ±8 sec around the poke. Compared to the averages around random time points, regions wide patterns of activity become apparent, with neurons located in different functional areas increasing and/or decreasing their firing rates with an order of succession that seems to hold over animals (given our sample size of three). A non-parametric statistical test (Maris and Oostenveld, 2007) between the poke and random firing rates reveals clusters over space and time that show a significant Monte-Carlo p-value (Figure 4, 3^rd^ column). These span most of the targeted regions and seem to mainly focus in time from 1 second before the poke to up until 6 seconds after. The test we used is designed to remove the effects of multiple comparisons but underplays any small but real effects. So we hypothesise that in experiments with larger sets of trials a more detailed pattern of firing rate modulations would emerge, maybe one that could be compared between subjects. Taking into account that the different regions targeted by the probe do not have any special anatomical connectivity (the cortical region targeted for example, i.e. the parietal cortex, is part of the association cortex while the targeted thalamic nuclei are mainly first order relays of the somatosensory system), the data resulting from our exploratory experimental design allow us to formulate the hypothesis that such patterns of activity might be ubiquitous in the brain with even more regions being part of this interplay of populational activity modulation.

**Figure 4.**
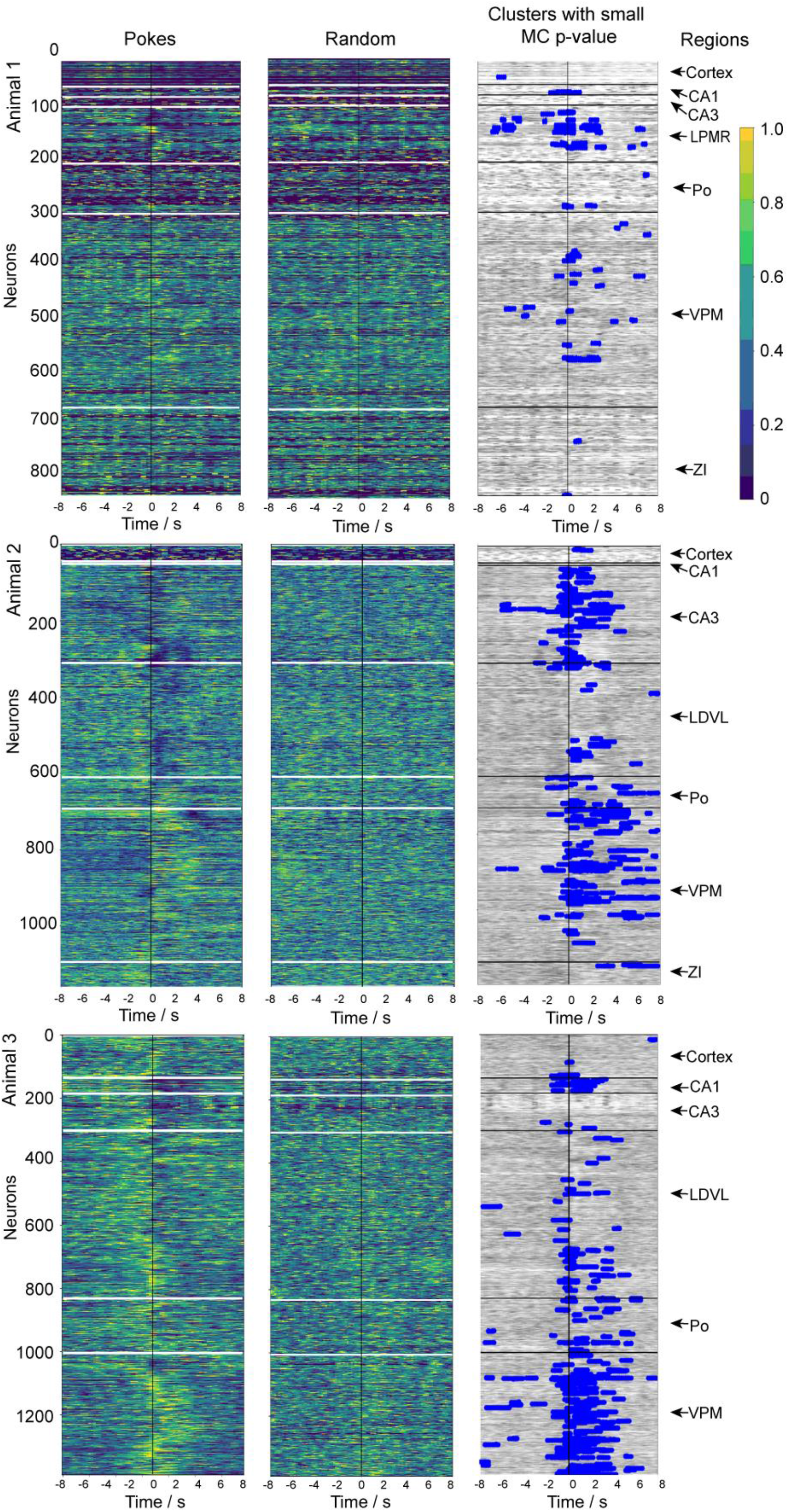
Global patterns of differential firing activity around specific events during a behavior task. The rows show the patterns formed in three separate animals (animals are the same as in Figure 2). Each figure in the first two columns shows the normalised firing rate of each neuron, averaged over trials and sorted in the y axis according to the depth of the neuron in the brain. For Animal 1 we averaged 83 trials, for Animals 2, 96 and for Animal 3, 100 trials. The first column shows the activity around the poking events and the second one around randomly selected time points over the duration of the recording (average of same number of random events as the number of trials averaged in column 1). The 3^rd^ column shows the results of a cluster based non-parametric statistical test between the poke and the random events data sets. The blue points (superimposed on the poke data set) are the clusters with a Monte-Carlo p-value smaller than 0.05 (the critical value for a one-sided t-test with alpha 0.05).

We argue that such global patterns have up to today gone unrecognised due to the limited number of regions standard electrophysiology probes can access. We also note how subcortical regions, not as easily accessible by optical methods, seem to play a prominent role in the brain’s generation of salient behavioural events. To test the hypothesis that multiple (even anatomical sparsely connected) regions play an important role in salient behavioural events one would need multi shank probes that can access multiple regions of the brain, both superficial and deep. Such probes have started to appear (Chung et al., 2019; Musk and Neuralink, 2019) and one can now use some of these technologies to target multiple regions anywhere in the brain. What these non-CMOS probe technologies though compromise on, in order to achieve recording spread, is electrode density.

### Density of local information correlates with electrode density

In order to test what a probe with large regional targeting but lower density would record in a comparable data set, we started from our chronically recorded data (day 10 for Animals 1 and 2 and day 7 for Animal 3) and decimated the recording electrodes, lowering the electrode density by a factor of 4 and of 16. These densities were chosen so that a 400 electrode probe would be able to target 10 mm with a single shank (factor of 4) and 4 shanks of 10 mm each (factor of 16). We then rerun the spike sorting analysis. A summary of how many neurons each electrode density design managed to detect can be seen in Figure 5 (with more details shown in Table 2). The reduced electrode density significantly diminished the quality of the local information gained defined by the number of total units recorded and by the number of single units extracted. Figure 5 shows that a reduction of density by a factor of 16 resulted in the reduction of all units detected by a factor of more than 6 and of the single units by a factor of more than 10.

**Figure 5.**
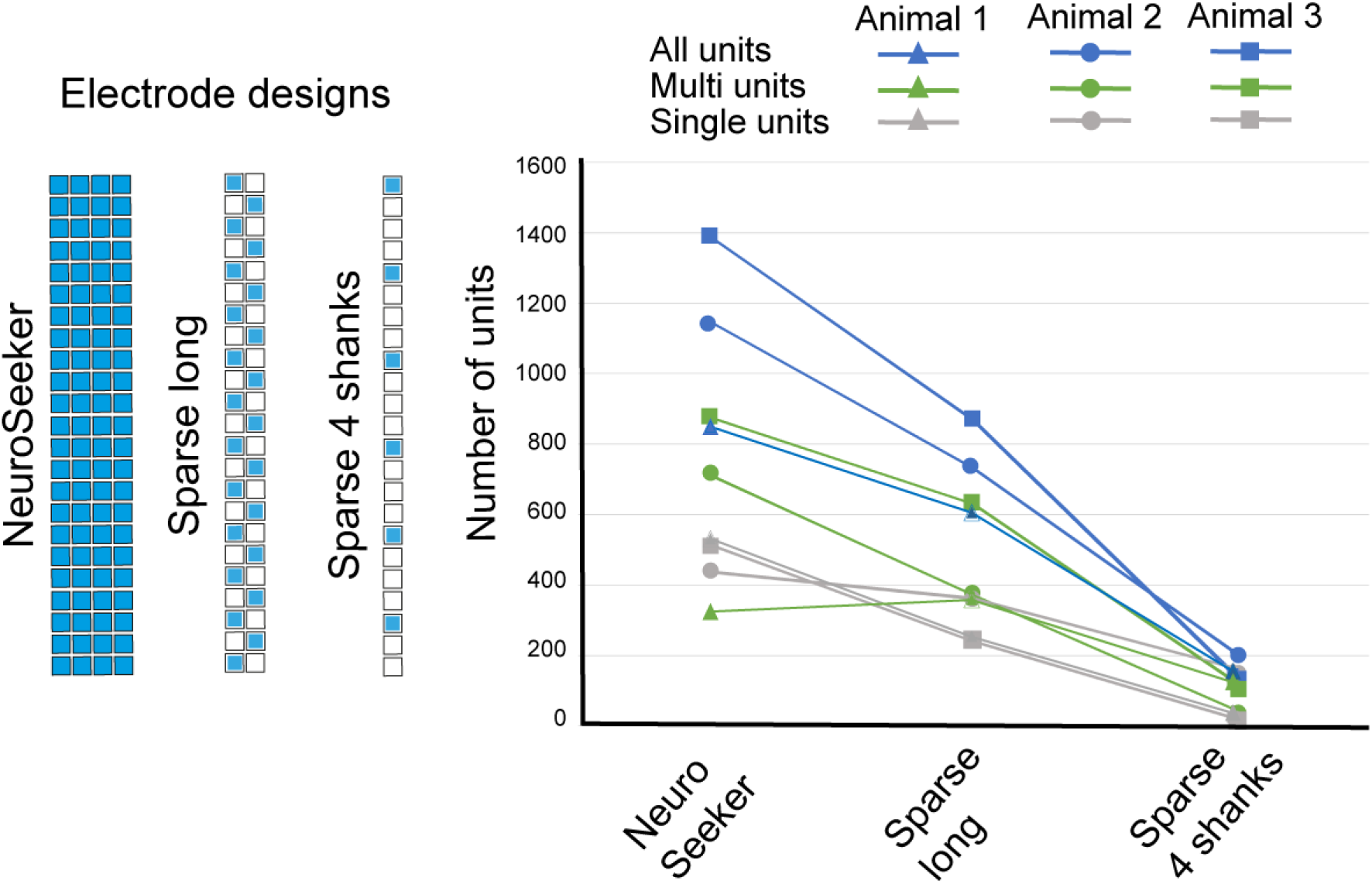
Different electrode densities and the drop in the number of single (green) and multi (grey) units (blue is the total units). The Sparse long configuration is a reduction of electrode density by 8 and the Sparse 4 shanks one by 16 compared to the original NeuroSeeker electrode design.

**Table 2.** Detailed results on the number of single, multi and total units detected in three animals with the different probe electrode densities.

| **Animal 1** | NeuroSeeker | Sparse Long | Sparse Long percentages | Sparce 4 Shanks | Sparce 4 Shanks percentages |
| --- | --- | --- | --- | --- | --- |
| Total units | 848 | 603 | 0.71 | 150 | 0.18 |
| Multi units | 320 | 354 | 1.11 | 119 | 0.37 |
| Single units | 528 | 249 | 0.47 | 31 | 0.06 |
| % multi units | 0.38 | 0.59 |  | 0.79 |  |
| % single units | 0.62 | 0.41 |  | 0.21 |  |

| **Animal 2** | NeuroSeeker | Sparse Long | Sparse Long percentages | Sparce 4 Shanks | Sparce 4 Shanks percentages |
| --- | --- | --- | --- | --- | --- |
| Total units | 1147 | 731 | 0.64 | 205 | 0.18 |
| Multi units | 436 | 361 | 0.83 | 162 | 0.37 |
| Single units | 711 | 370 | 0.52 | 43 | 0.06 |
| % multi units | 0.38 | 0.49 |  | 0.79 |  |
| % single units | 0.62 | 0.51 |  | 0.21 |  |
| **Animal 3** | NeuroSeeker | Sparse Long | Sparse Long percentages | Sparce 4 Shanks | Sparce 4 Shanks percentages |
| Total units | 1389 | 868 | 0.62 | 140 | 0.10 |
| Multi units | 876 | 629 | 0.72 | 121 | 0.14 |
| Single units | 513 | 239 | 0.47 | 19 | 0.04 |
| % multi units | 0.63 | 0.72 |  | 0.86 |  |
| % single units | 0.37 | 0.28 |  | 0.14 |  |

### Detailed spiking activity over many regions correlates with patterns of complex behaviour

We then asked if the extra information arising from the detailed knowledge of single neuron activity would be beneficial in defining brain states that corresponded more accurately with behaviour outcomes. In order to create a preliminary test for this we first defined behavioural states as follows.

We cropped the behavioural video around the animal and then subsampled it by averaging every 12 frames (creating a new video of 10 fps). We then run the cropped and subsampled video through a principal component analysis (PCA) algorithm and kept the top 100 components. Figure 6A shows in four inserts four average frames and their equivalent reconstructions from the 100 top PCs of the day 10 of Animal 2 recording. We then passed these 100 PCs of all frames to the t-SNE algorithm (Dimitriadis et al., 2018) and finally used DBSCAN to cluster the resulting 2D embedding. Figure 6A shows the results of this process. Each cluster corresponds to a specific combination of the animal’s place in the arena and its body posture. The arena had enough unique landmarks that one can easily identify the animal’s position in it even in the cropped video. For example, clusters blue and red in Figure 6 are both defined by the times that the animal is at the poke with its body angled at about 120 degrees off the horizontal, but differ by the fact that in the red cluster the animal is rearing so its body shape is more spherical.

**Figure 6.**
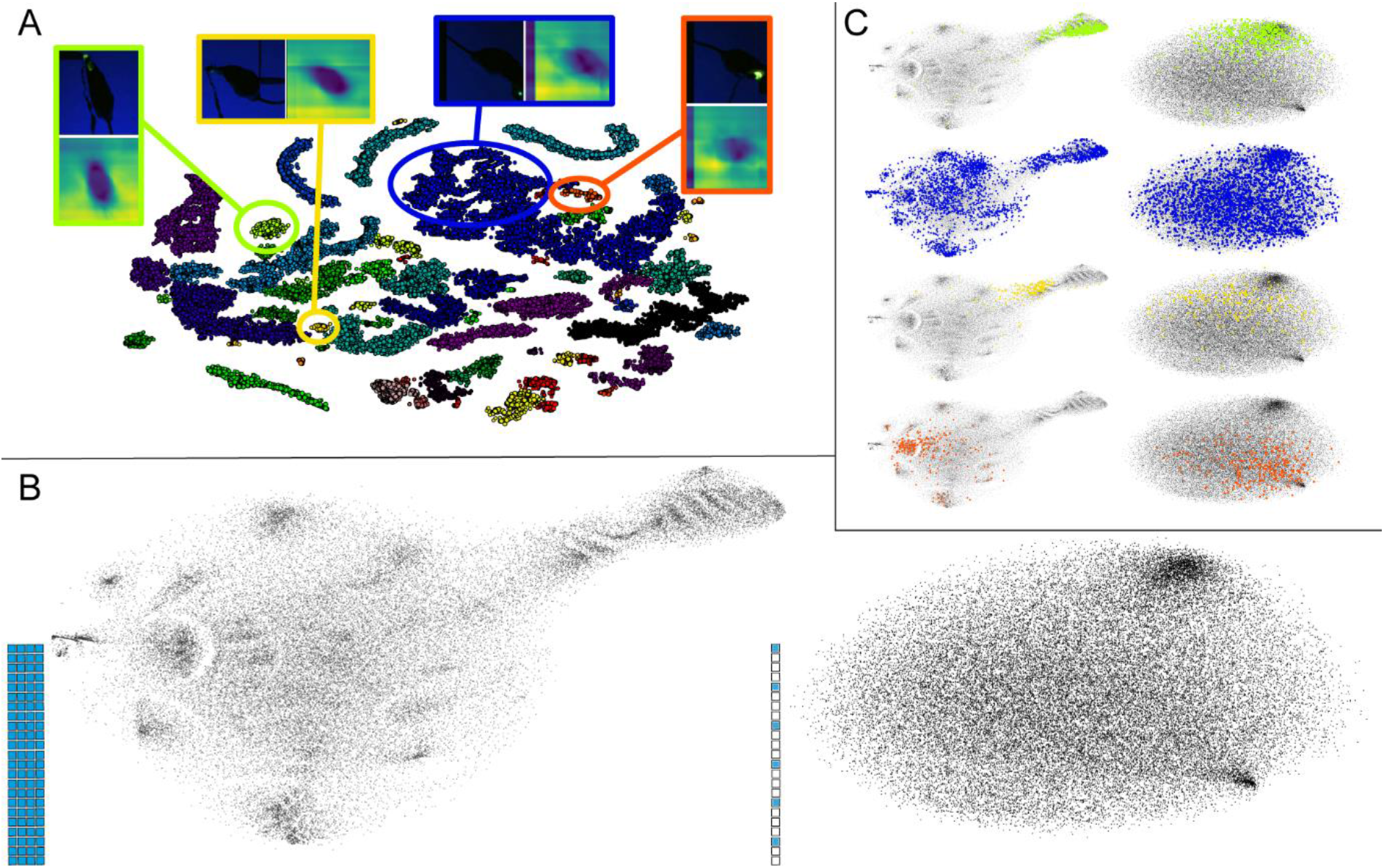
The reduction in the density of the recording electrodes is followed by a reduction in the detection of self-similar brain activity patterns. A) t-SNE dimensionality reduction and DBSCAN clustering results of the cropped and averaged behavioural video. The 4 inserts show a characteristic example of the frames in the 4 marked clusters. The top/left of the insert is the actual frame and the bottom/right the reconstruction of the top 100 PCs (which is what was fed into the t-SNE algorithm). B) t-SNE embedding of the NeuroSeeker (left) and the Sparse 4 shanks (right) recording (firing rates of all neurons) of that session. C) Same t-SNEs as in B but with the points that clustered in the four clusters selected in A being superimposed.

Having defined a set of low level behavioural states (described by position and posture) we then followed the same process (PCA and t-SNE) to 2D embed the average firing rates per 100 ms for all neurons. We did this for both the NeuroSeeker density and for the Sparse 4 shanks one (16 times sparser). The results of this embeddings is shown in Figure 6B for Animal 2. In the case of the NeuroSeeker probe, a large percentage of the firing rate vectors aggregated in self similar regions. In contrast, the Sparse 4 shanks embedding, shows that a drop in single neuron spiking information makes it impossible to differentiate between unique brain states in an unsupervised learning fashion. The same loss of self-similar clustering due to the lowering of the density of the probe is shown in the other two animals we recorded from (seen in Supplementary Figures 2B and 3B)

Each point in the embedding of the behaviour video has a corresponding point in the embeddings of the two firing rate embeddings (corresponding to the same 100 ms window). The question we wanted to answer was whether the points that cluster together in the behavioural video would have correspondence to local regions in the brain activity embeddings.

For Animal 2 out of the 72 behavioural clusters defined, 66 had no obvious correspondence to sub regions in the brain activity embedding (example of that is the blue cluster in Figure 6C). On the other hand, 6 of them showed a very localised pattern on the NeuroSeeker brain activity embedding, where similar behaviour corresponded to highly self-similar neural patterns (see Figure 5C showing three of these 6 clusters (yellow, red and green)). The second column of Figure 5C shows that this was not the case for the Sparse 4 shanks embedding where these 6 behavioural clusters (three shown but the other three behaved similarly) showed a disperse pattern on the t-SNE embedding. So a drop in density, resulting in a drop in single neuron spiking information also resulted in the loss of the few brain activity regions whose self-similarity corresponded to self-similar behaviour.

Similar results were obtained from the two other animals (see Supplementary Figures 2 and 3) where the behavioural clusters showing a high degree of self-similarity in the corresponding firing rate patterns stopped doing so when collected by a smaller density probe. We quantified this loss of self-similarity of the brain activity patterns over the different behaviourally defined clusters by calculating the average distance of points on the firing rate t-SNE for each one of these clusters and then aggregating the results over all clusters for each recording (animal / probe type). The results of this metric can be seen in Supplementary Figure 4 showing that the groups of points of the firing rate t-SNE corresponding to the behavioural clusters show a smaller average spread in the NeuroSeeker recordings vs. the Sparse 4 shanks ones.

### Do we need all these electrodes?

Integrating CMOS-based scanning technology into the shaft of an *in* vivo neural probe has now made it possible to drastically increase electrode density. However, given the challenges associated with the volume of data generated by such probes, neuroscientists must decide whether further increases in recording density are actually useful. Current approaches to spike sorting would appear to benefit from higher density probes (Dimitriadis et al., 2018; Moore-Kochlacs, 2016; Rossant et al., 2015). Furthermore, initial results (Jun et al., 2017a) show that increased electrode densities could help compensate for drift in chronic recordings and allow following individual neurons over the course of days and weeks. Finally, CMOS technology enables the fabrication of electrodes much smaller than a neuron’s soma and dense enough to capture fine details of a single neuron’s extracellular field throughout its dendritic tree (Delgado Ruz and Schultz, 2014; Jia et al., 2018).

Whether useful information is present at these scales, and whether it can contribute to the further understanding of brain function, is an open question. Thus, in order to address whether ultra-high density in neural recordings will be useful, we collected a series of datasets with an ultra-dense probe during anesthesia from many different brain regions (Figure 7A). These data sets are available online (http://www.kampff-lab.org/ultra-dense-survey/) and a summary of them is given in Supplementary Table 2.

**Figure 7.**
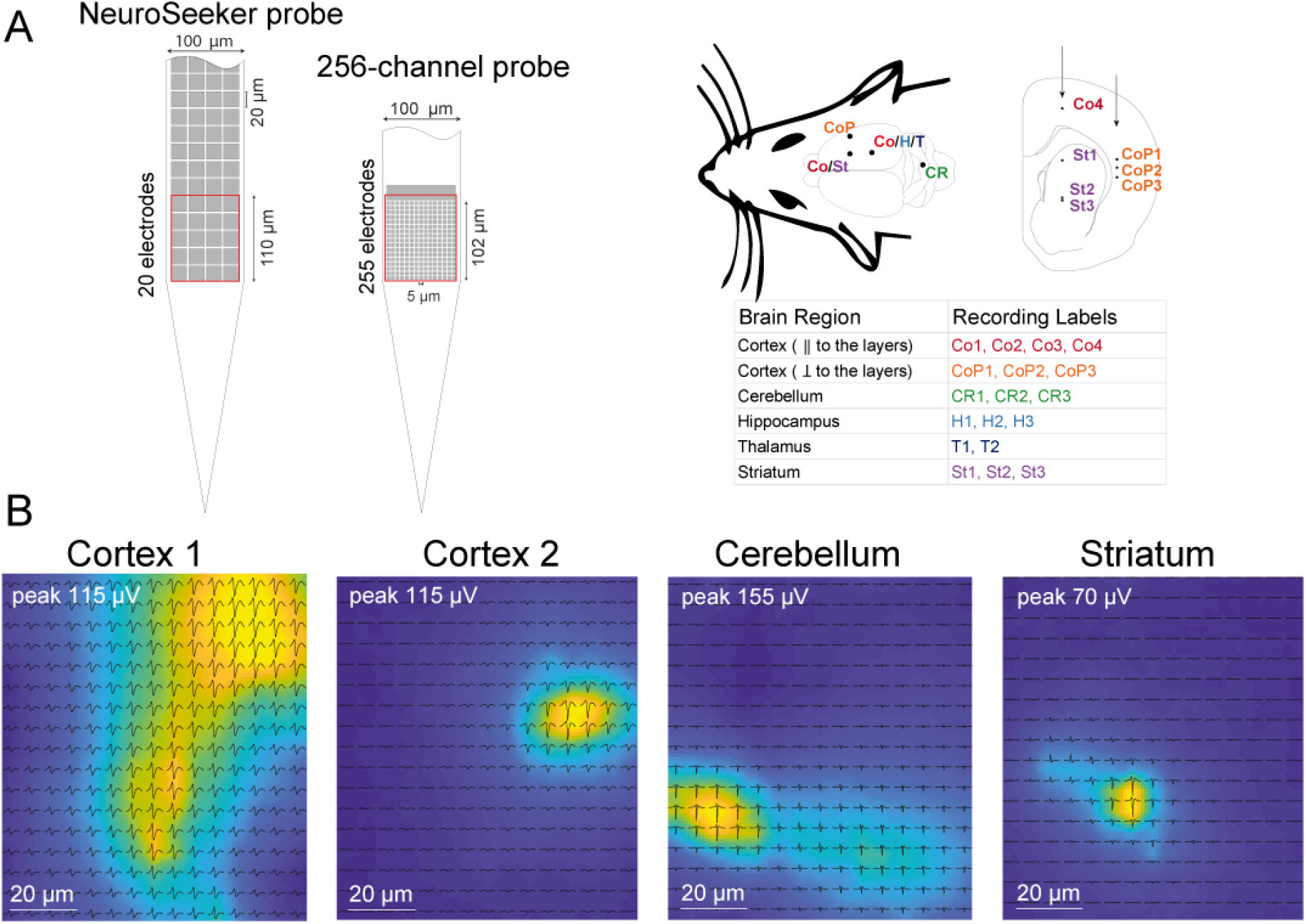
A) The 256-channel ultra-dense probe packs 255 (15 × 17) electrodes in the same area (100 × 102 μm) as 20 electrodes from the NeuroSeeker CMOS scanning probe representing a 13-fold increase in electrode density. We conducted four separate penetrations covering five anatomically distinct brain regions and two different orientations (i.e., parallel and perpendicular) with respect to the laminar architecture of the cortex, leading to 18 separate recordings. B) Four example neurons recorded with the ultra-dense probe (the cortical ones were from Co recordings). For each neuron, the mean waveform across spikes on each site is shown in black, arranged spatially according to recording site locations. The colormap depicts the amplitude of the spike at each location in space, taken from the mean waveforms then spatially up-sampled and interpolated. Sites with abnormally large impedance (see text) were excluded and waveforms at those locations were estimated by interpolation. The amplitude of the mean waveform on the largest channel, corresponding to the max color value, is given as the ‘peak’.

The ultra-dense probe contains 255 electrodes (and one large reference) with an area of 5 × 5 μm and separated by 1 μm, and was fabricated by imec as part of the NeuroSeeker project (see Supplementary Table 3 for a review of the CMOS and ultra-dense passive probes). As shown in Supplementary Figure 5, the probe electrodes have a low impedance value at 1 kHz of 790 ± 100 kΩ. Note that 16 of the 255 electrodes are non-functional because their impedance value at 1kHz is higher than 2 MΩ. The noise magnitude computed in saline for the functional electrodes is 5.09 ± 0.03 μV.

The data shown in Figure 7B illustrate the ability of this ultra-high-density array of electrodes to detect the activity from the same neuron on several adjacent electrodes thus generating a sort of electrical image of the neuron. This preliminary analysis shows the potential of such densities for the further study of the correlations of this electrical profile with a neuron’s position in the brain, its shape or its electrochemical properties. More importantly one can now envision a possible future CMOS probe combining the same resolution as the 255 channels probe with a recording area equal to the NeuroSeeker probe (which would roughly result in a 20,000 channel probe). The operational implications of such technology are admittedly intimidating (20K channels means almost 3TB of data per hour). Yet, connecting single cell morphology and detailed, sub-cellularly resolved, electrical behaviour with network activity from many regions of the brain is a capability the field should consider exploring in more depth (since the technology is finally here).

## Discussion

The NeuroSeeker project is the culmination of a number of efforts to translate major advances in microfabrication into new tools for electrophysiology (Neves et al., 2008b; Torfs et al., 2011; Lopez et al., 2013; Jun et al., 2017b; Raducanu et al., 2017; Herbawi et al., 2018; Dorigo et al., 2018; Seidl et al., 2011). The project has produced a device with multiplexing electrode scanning circuitry within the thin shaft of an implantable neural probe, which provides the highest number of simultaneous outputs for a single shaft *in vivo* extracellular recording device ever achieved.

Here we report that the integrated circuitry of the NeuroSeeker probe does not have an adverse effect on its ability to record neural activity during both acute and chronic recordings. We show that when compared to a passive probe with the same electrode configuration, the NeuroSeeker probe offers simultaneous recordings from >1000 electrodes with similar signal-to-noise ratio. We also show that the probe can be used in long term chronic experiments with a signal deterioration over time comparable to other silicon based probes. Finally, we demonstrate that the probe is both mechanically and electrically robust enough to be removed at the end of a chronic experiment and be reused without any loss of signal quality. Given all this, we propose that integrating on-shank scanning circuitry is not only a promising technology for creating ultra-high density *in vivo* probes, but will, at the very least, allow the simultaneous recording from the entire probe with a density sufficient for accurate spike sorting.

The promise of such high-density, large electrode count probe designs, with their capability to scale to even larger numbers and densities, offers more than simply increasing spike sorting accuracy; they make it possible to simultaneously extract information from the brain on three different scales. At the single neuron level, high densities provide valuable information about cell type and signal propagation throughout the dendritic tree, which can help address the underlying single cell computation. At the level of the local circuit, recording hundreds of neurons simultaneously from a given functional region (here we record from more than 200 per region) is required to test many major hypotheses in systems neuroscience, e.g. the canonical circuit, the engram, or Hebbian learning. At the level of the global network, recording from a number of regions throughout the brain allows studies of how the brain as a large network of functionally and anatomically diverse regions generates complex behavior. A combination of these two capabilities, we argue, is also indispensable for executing exploratory experiments, designed not to answer a specific hypothesis but to rather generate a range of new ones.

The results we present here demonstrate how the probe was utilised to formalise such new hypothesis. Using its global scale capabilities, a new view of brain inter-regional communication during salient behavioural points (where the animal acts in order to update its model of the world) was demonstrated. This view allowed us to hypothesise that there exists a large brain network composed of many and anatomically sparsely connected regions that both activates before the behavioural event and subsequently computes its results. Focusing on the probe’s detailed local scale information we hypothesise that one can extract correspondences between data defined epochs extricated from complex behaviour on one hand and patterns of firing rates on the other, as long as one has at their disposal a rich enough view of the brain’s dynamics. It is exactly such correspondences that can provide the strong background onto which theories like the canonical circuit, the engram or reinforcement learning can be tested and implementation details of their algorithms can be formulated.

The CMOS technology makes future devices, which distribute dense electrode groups throughout the brain possible. Our results indicate that their exploration can greatly benefit neuroscientific research, that being also one of the major aims of large neuroscience projects like the NIH Brain Initiative (Koroshetz et al., 2018).

The number of simultaneously recorded electrodes of the NeuroSeeker probe created new challenges for previously simple aspects of an experiment, e.g. online data monitoring. We also presented here a GPU-based method for generating a real-time visualization of the very large data sets generated by these probes. This greatly facilitates the control of an ongoing experiment and provides an intuitive overview of large scale structures present in the spiking activity recorded by the probe.

Finally, the high costs associated with CMOS development necessitate a large initial investment to produce just a small number of prototype devices. However, the promise of modern CMOS production scaling is that, after an expensive development phase, a large number of equivalent devices can be produced at a tiny fraction of the initial costs. The recent CMOS-based neural probe prototyping efforts, such as NeuroSeeker, have resulted in truly revolutionary devices for neurophysiology. However, a true revolution in neuroscience will only come when these devices are available in large numbers, to the entire community, for the price of a spool of tungsten wires.

## Methods

### Animal surgeries

All experiments were conducted with Lister Hooded rats ranging between 400 g to 700 g of both sexes. For both acute and chronic surgeries, the animals were anesthetized, placed in a stereotaxic frame and undergone a surgical procedure to remove the skin and expose the skull above the targeted brain region. For the acute surgeries we used urethane (1.6 g/kg, IP) anesthesia, while for the chronic ones we used isoflurane gas anesthesia (2% v/v). At the initial stage of each surgery the animal was also injected with atropine (0.05 mg/kg), temgesic (20 μg/kg, SC) and rimadyl (5 mg/kg, SC). Small craniotomies (about 1 mm in diameter) were performed above the target area. During the surgeries equipment for monitoring the animals’ body temperature and a video system for guiding probe insertion (Neto et al., 2016) was used. The targeted insertion coordinates were scaled according to the size of each animal’s skull. We measured the distance between bregma (B) and interaural line (IA) for all animals to find the ratio between the animal skull and the reference skull (B – IA = 9 mm) generated as part of the 3D atlas described in (Dimitriadis et al., 2014). Finally, we adapted the insertion coordinates for targeting specific brain regions with the use of the 3D atlas.

The insertion was done using an Angle Two Small Animal Stereotaxic Instrument (Leica Biosystems, US) which had its z-axis motorised (in house development) allowing precise control of the step size and speed of insertion. In two animals (Animals 1 and 3) the speed of insertion was 133 μm / min while for one animal (Animal 2) it was 1600 μm / min. In all cases the insertion step was 6 μm and the speed was controlled by the duration between steps.

Animal experiments were approved by the local ethical review committee and conducted in accordance with Home Office personal and project (I19EFD9C9; 70/8116) licenses under the UK Animals (Scientific Procedures) 1986 Act.

### Recordings

The CMOS-based probes are assembled on a PCB where the reference (REF) and ground (GND) wires are connected. The recording system consists of a headstage, which configures and calibrates the probe and serializes probe data, a microcoax cable and the base station board with a deserializer chip which connects to a commercial FPGA development board (Xilinx KC705). The computer used for controlling the NeuroSeeker system is connected to the FPGA board via a 1000 kBps-T Ethernet card. This recording system together with the FPGA code and low level C drivers for FPGA - computer communication were developed by imec. The FPGA code and the C drivers remain closed source. The protocol for the communication with the low level C drivers for the acquisition and saving of the data and subsequent visualization was implemented in Bonsai. Both Bonsai and the specific NeuroSeeker communication protocol are open source (Bonsai, 2017; Lopes et al., 2015). The CMOS extracellular signals were sampled at 20 kHz with a 10-bit resolution.

Due to the small diameter of the microcoax cable and the size of the behaviour arena there was no requirement for the use of a commutator during the freely moving animal recordings. No cable tension or significant torsion ever developed during the 1-1.5 hour long experimental sessions. The behaviour arena is a 1 × 1 m box with a back projection glass surface as floor onto which a computer projects imagery at a frame rate of 120 Hz. A grasshopper 3 (FLIR) camera is recording the movements of the rat in the arena, also at 120 Hz, while the controlling computer system is live tracking the animal generating x, y positions for each frame.

For the 256-channel probe recordings we used the Open Ephys (http://www.open-ephys.org/) acquisition board along with two RHD2000 128-channel amplifier boards that amplify and digitally multiplex the extracellular electrodes (Intan Technologies). Extracellular signals in a frequency band of 0.1–7,500 Hz were sampled at 20 kHz with 16-bit resolution and were saved in a raw binary format for subsequent offline analysis using Bonsai.

### Impedance measurements

For the 256-channel probe, impedance tests (at 1 kHz) were performed using a protocol implemented by the RHD2000 series chip (Intan Technologies) with the electrodes placed in a dish with sterile PBS, 1 mM, pH 7.4 and a reference electrode, Ag-AgCl wire (Science Products GmbH, E- 255).

### Probe removal

The probe removal from a chronic implantation starts by aligning the rat’s head (after euthanasia) with the stereotactic coordinates system. This was done by making sure the probe holder’s top surface is parallel to the Anterior-Posterior (AP) / Medial-Lateral (ML) plane of the stereotaxic frame. To achieve this, we make sure this plane is perpendicular to gravity using a spirit level. We then make sure the same thing is true for the rat’s head in the stereotaxic frame, i.e. the top surface of the probe’s holder is also perpendicular to gravity (using again a spirit level). This step follows from the fact that during probe implantation the probe was held by the stereotaxic frame (while glued on the rat’s skull) in such a way that it’s top surface was parallel to the frame’s AP/ML plane, so by realigning those two planes we bring the probe back to the original position it was implanted in respect to the Dorsal-Ventral (DV) movement of the stereotaxic frame’s arm. After alignment, we connect the probe holder to the frame’s arm, and then cut through the thin layer of resin (that is kept away from the probe by the tapered skirt), thus separating the part that stays attached onto the head and the part that holds the probe. Then we slowly remove the detached probe by withdrawing the stereotaxic frame’s arm. The alignment of the head and the connection of the arm to the still attached probe holder is of importance since any tension between the head and the frame’s arm due to misalignment will be released after the cutting of the resin in the form of a movement of the head in respect to the arm resulting in the breaking of the probe shaft. After removal a series of trypsin and ddH_2_O cleaning cycles were undertaken until, under a microscope, no organic or salt residues could be seen on the probe’s shaft.

### Analysis

For the analysis, the CMOS-based probe recordings were filtered with a band-pass of 500-3,500 Hz and saved in binary format using a Bonsai interface. When the external reference was selected, to diminish the effect of artefacts shared across all channels, we subtracted the median signal within each group across the recording electrodes from the respective group.

For the reconstruction of a unit’s position on the probe (for the results shown in Figure 3) we used a weighted (according to peak to peak amplitude (PPA)) average of the positions of the electrodes that showed a significant deviation from baseline on the average spike waveform. That is, we selected the electrode with the highest PPA at the time around 0 of a unit’s spike template, found all other electrodes whose PPA was over a threshold and then moved the position of the unit from the highest electrode’s position towards the other relevant electrodes in a linearly weighted manner according to their PPAs.

For the firing rate activity time locked to an event (exploratory pokes or randomly selected time points) we first calculated the firing rates of all neurons by summing all of the spikes of each neuron found in a bin of 8.3 ms (equal to one frame of the 120 fps camera we have been using to record the animals’ behavior). We then further averaged these firing rates with a moving window of 30 (giving firing rates of 250 ms resolution). We then took 32 bins before and 32 bins after each event (±8 sec) and averaged those firing rate time series together. For Animal 1 there were 83 poke events, for Animal 2 96 and for Animal 3 100 events. We used the same number of events to create the average around random time points. The resulting event time locked firing rates were then normalised individually to range between 0 and 1.

To test whether the firing rate of the poke averages differed from the ones of the random averages we used a non-parametric cluster based test as described in (Maris and Oostenveld, 2007). We used a one-sided test with an alpha-level of 0.05 (95^th^ percentile) for the t-test used to select samples for clustering and a Monte-Carlo alpha-level of 0.05 for calculating the p-value of the perturbation distribution. The minimum area (number of points) for a cluster was set to 5.

For the t-SNE embeddings of the firing rates, we first created average firing rates as above but with a 100 ms window (averaging 12 frames). We then run PCA on the number-of-electrodes X number-of-windows matrix and kept the top 40 principal components (in Animal 2 for example this described 13% of the NeuroSeeker’s data variance and 33% of the Sparse 4 shank’s one with similar percentages for the other two animals). We finally used t-SNE to embed them in a 2D space. This was run with a perplexity of 100, a theta of 0.3, an eta of 200, exaggeration of 12 and 6000 iterations.

For the video embedding and clustering we first cropped the video around the rat (cropped frame size = 250 × 250 pixels), then averaged every 12 frames (creating the same 100 ms window per data point as above), rescaled to 80 × 80 pixels, greyscaled and flattened. The resulting 6400 X number-of-time-windows matrix of frames was then run through PCA. We kept the top 100 PCs (95% explained variance in all animals) and put those through a t-SNE algorithm with the same settings as above. The DBSCAN algorithm that was run on the resulting embedding had an epsilon of 0.025 and a minimum samples of 40.

To calculate the drop in self-similar aggregation of the firing rate vectors on the firing rate t-SNE with the drop of density we first created the average distance (Euclidian) on this embedding of 20000 randomly selected points. Then for each group of points that corresponded to a behavioural t-SNE / DBSCAN cluster (i.e. points that clustered together in the behavioural t-SNE) we calculated their mean distance on the firing rate t-SNE. This mean distance we then divided with the average distance of the entire t-SNE calculated before. This gave us, for each recording of interest, a number of average distance ratios, one for each behavioural cluster, that measured how much the group of points belonging to that cluster aggregated on the firing rates t-SNE. We then calculated the mean and 95^th^ percentile confidence intervals (using a 10000 iterations perturbation statistic) over all clusters of this ratio for each recording of interest.

For the 256-channel probes, a third order Butterworth filter with a band-pass of 250-9,500 Hz (95% of the Nyquist frequency) was used in the forward-backward mode. The noise magnitude was computed as an estimate of the background noise, *σ_Median_* = median(| signal(t)|/0.6745), of each filtered voltage-time trace during 5 seconds. Some results were represented as mean ± standard deviation.

## Acknowledgments

Joana Neto was supported by the fellowship SFRH/BD/76004/2011 from Fundação para a Ciência e Tecnologia, Portugal.

Susu Chen was funded by the Wellcome Trust as a Sir Henry Wellcome postdoctoral fellow.

Richárd Fiáth is thankful to the Hungarian National Research, Development and Innovation Office (PD124175).

This work was supported by funding from the European Union’s Seventh Framework Programme (FP7/2007-2013) under grant agreement nr. 600925 (NeuroSeeker), by the Bial Foundation (Grant 190/12) and by the Hungarian National Brain Research Program 2017_1.2.1-NKP-2017-00002.

## Notes

#### Summary of Updates

Added more animals results for the analysis presented

