## Supplementary Material for "Why not record from *every* electrode with a CMOS scanning probe?"

Supplementary Information

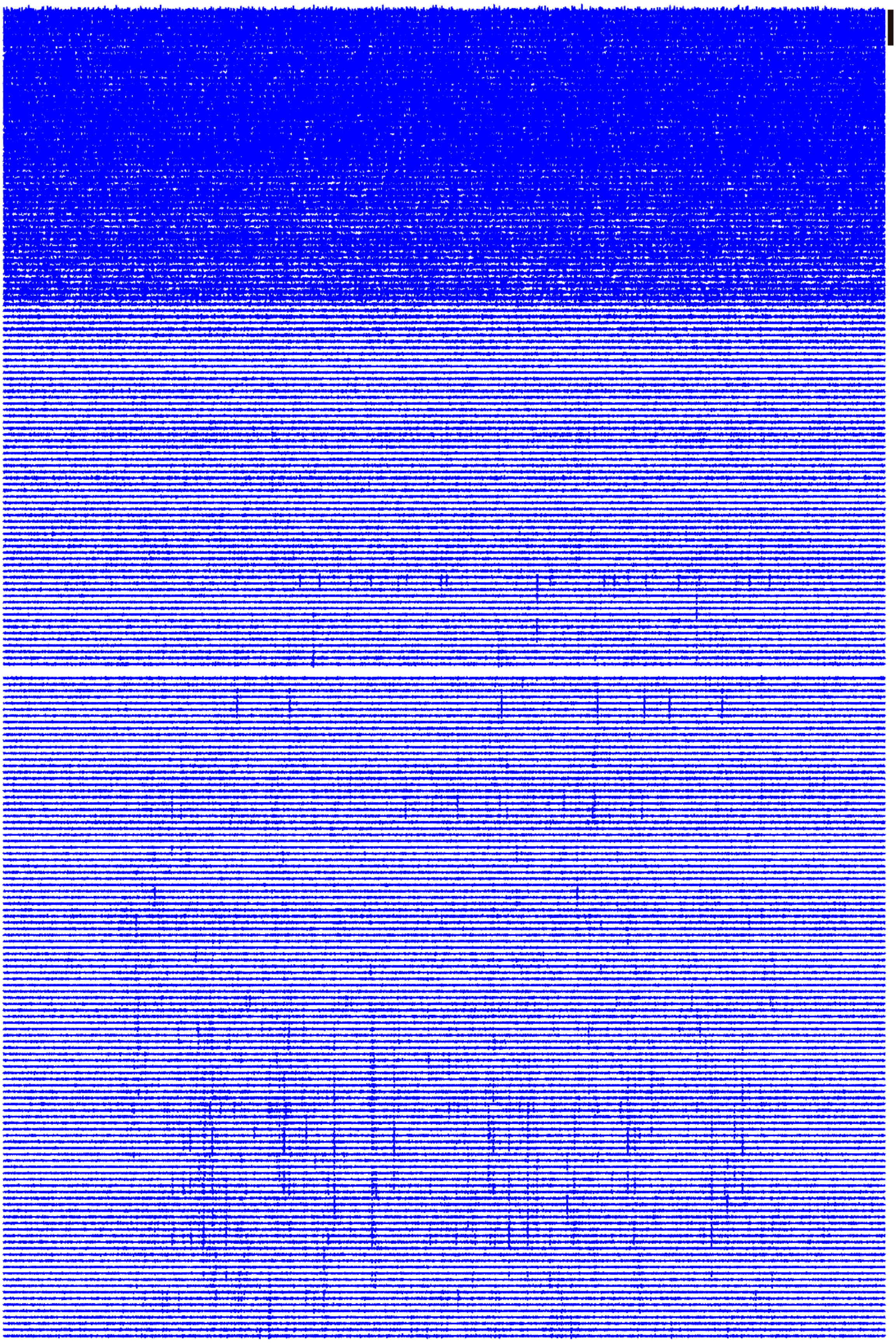

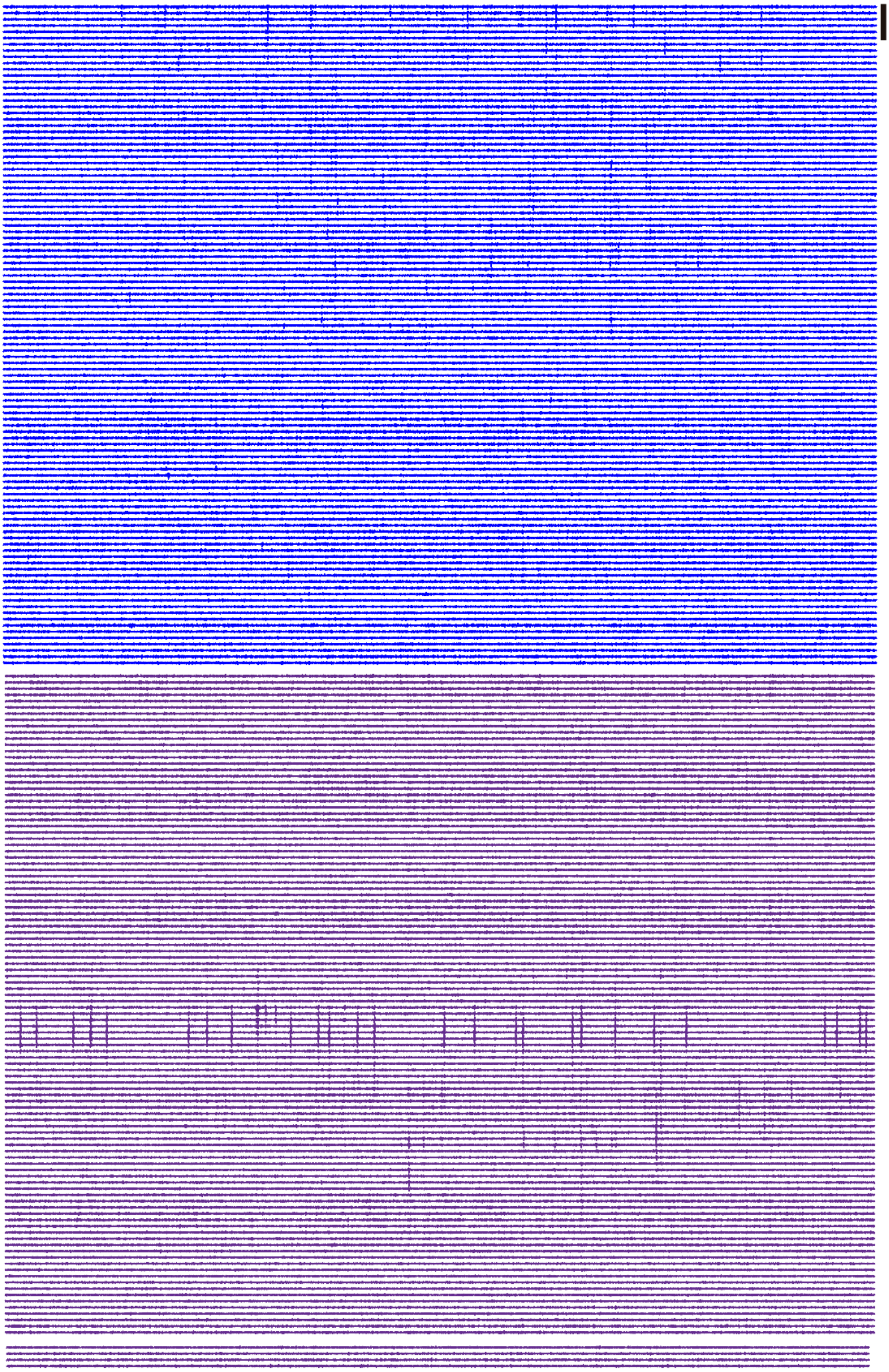

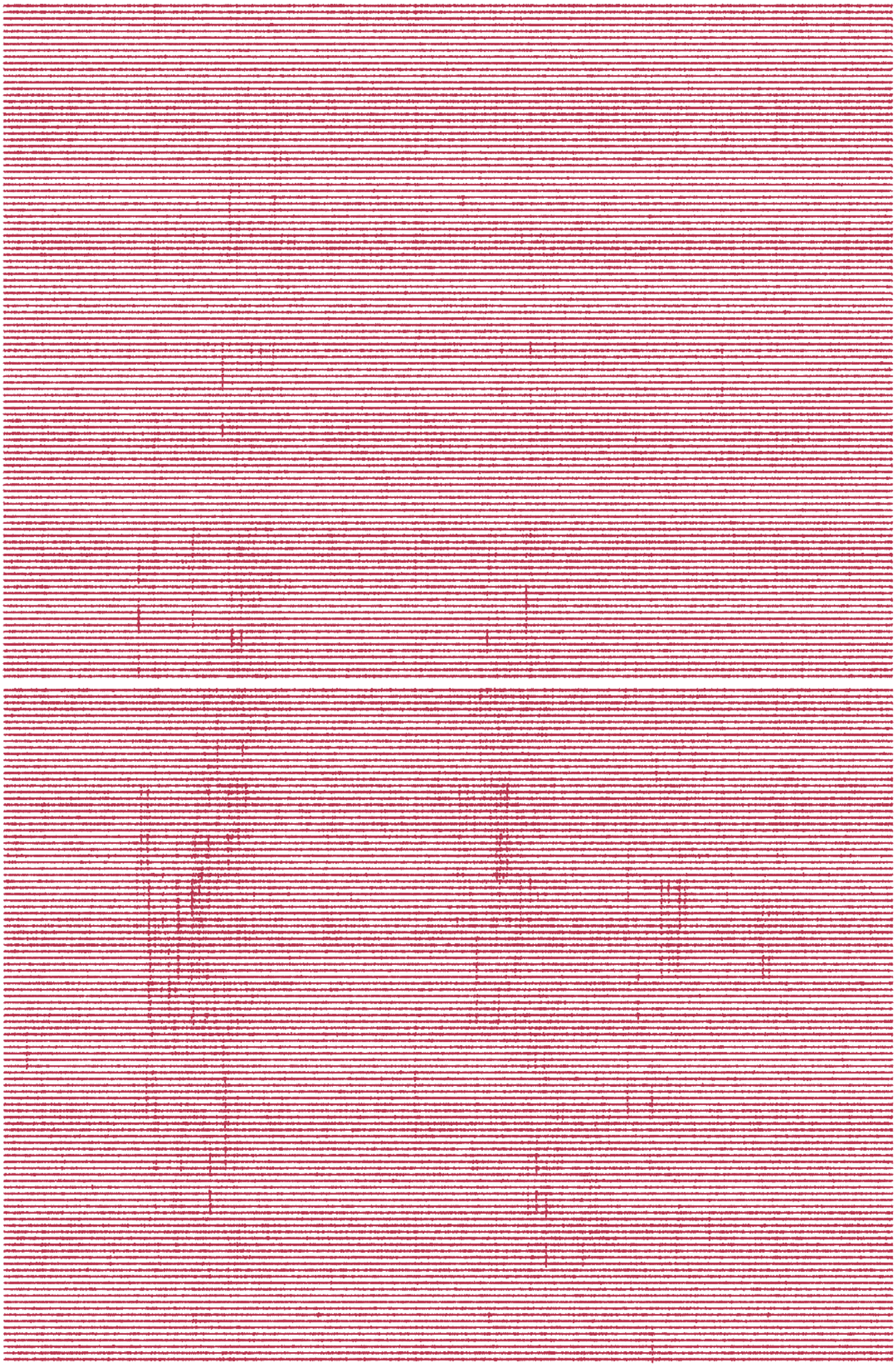

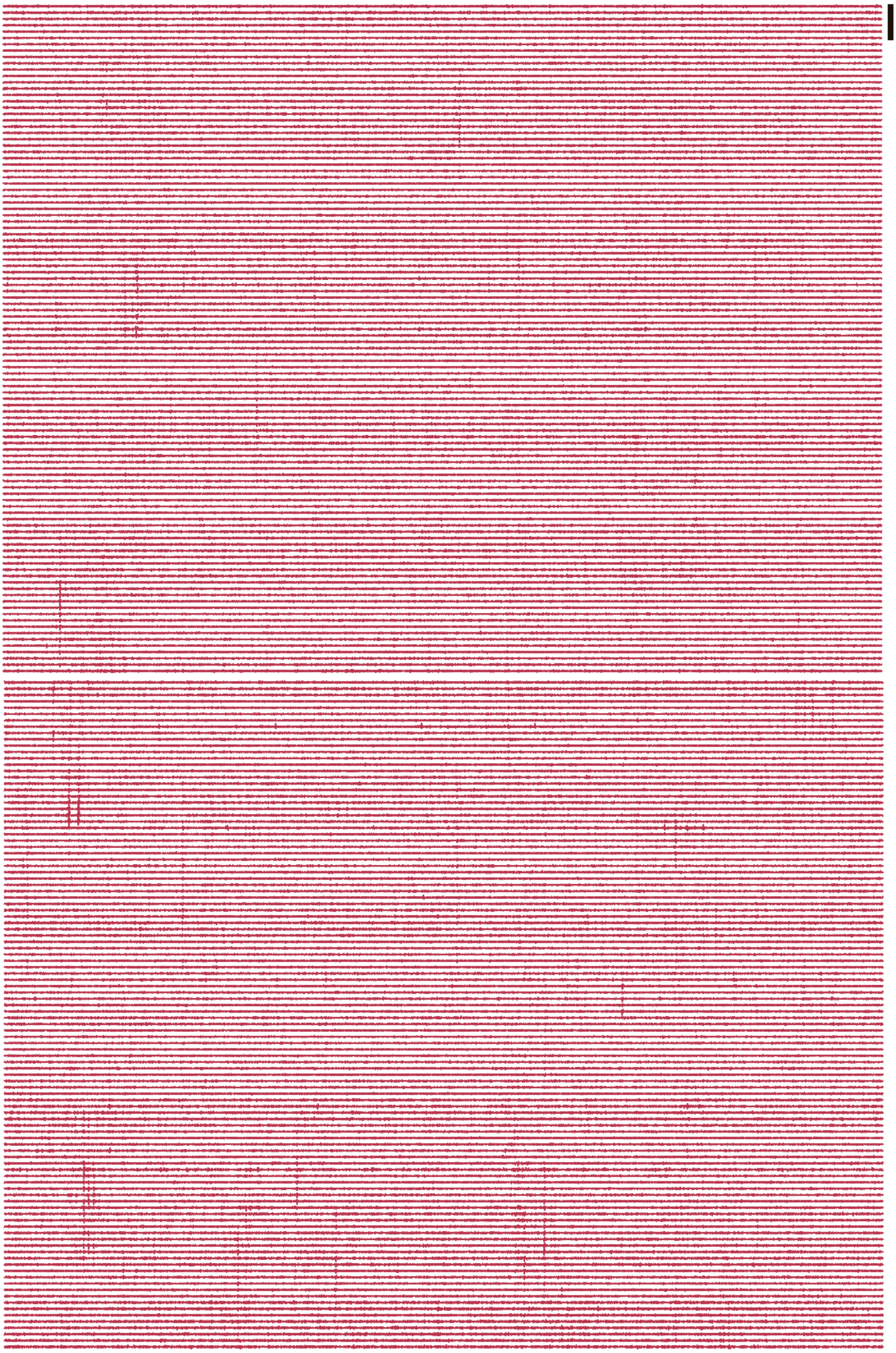

Figure 1. Example of a recording (*18_26_30.bin*) segment performed by a CMOS scanning neural probe with 1060 electrodes set to AP mode. 500-ms-long AP traces from a probe spanning multiple brain regions (cortex shown in blue, hippocampus shown in purple and thalamus shown in red). In each page, two groups from the probe are represented; scale bar= 600 µV.

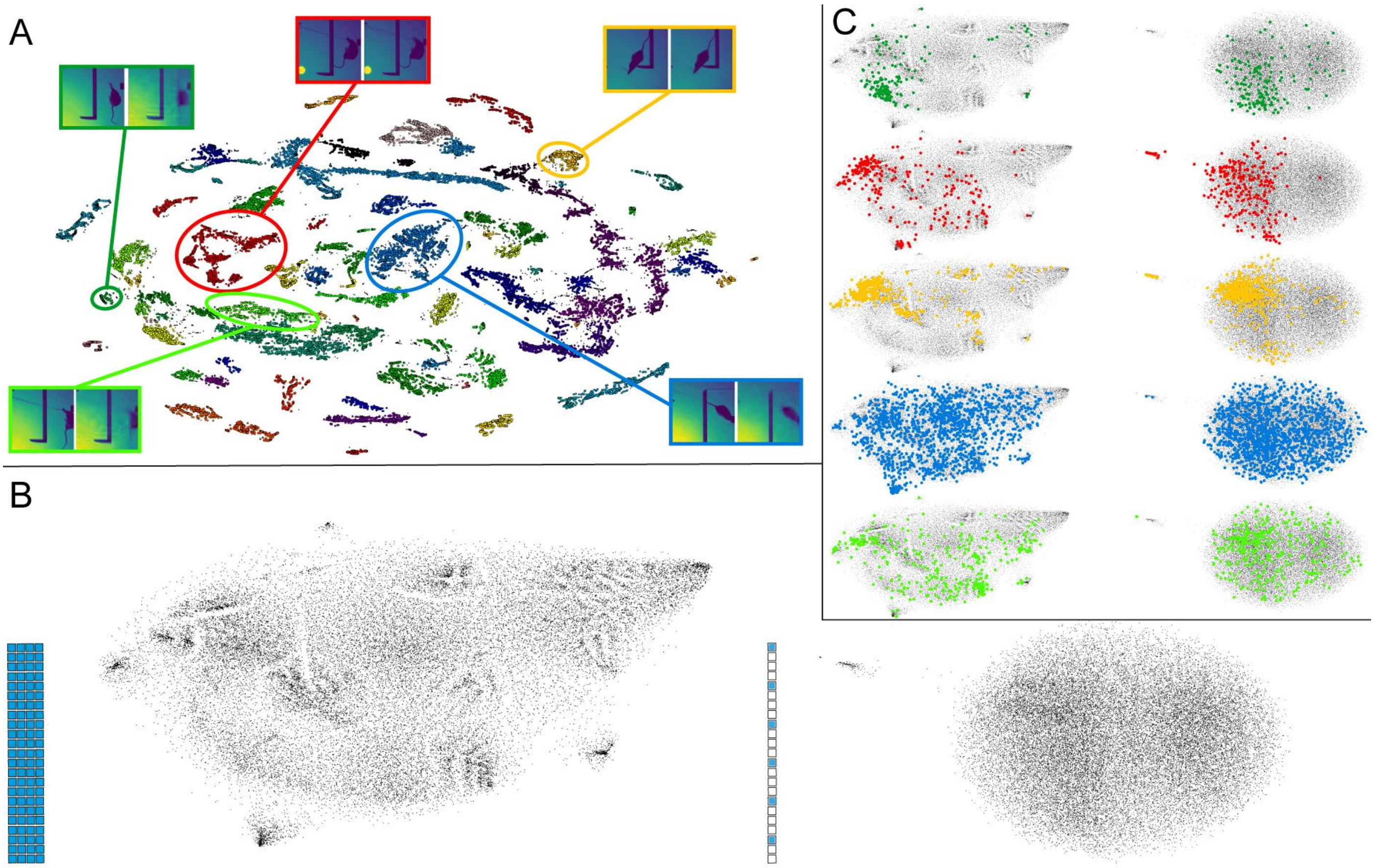


Figure 2. Loss of correspondence between self-similarity of behavior and of the firing rates of recorded neurons with a decrease of the probe's density (see Figure 6) for Animal 1. A, B and C identical to Figure 6.

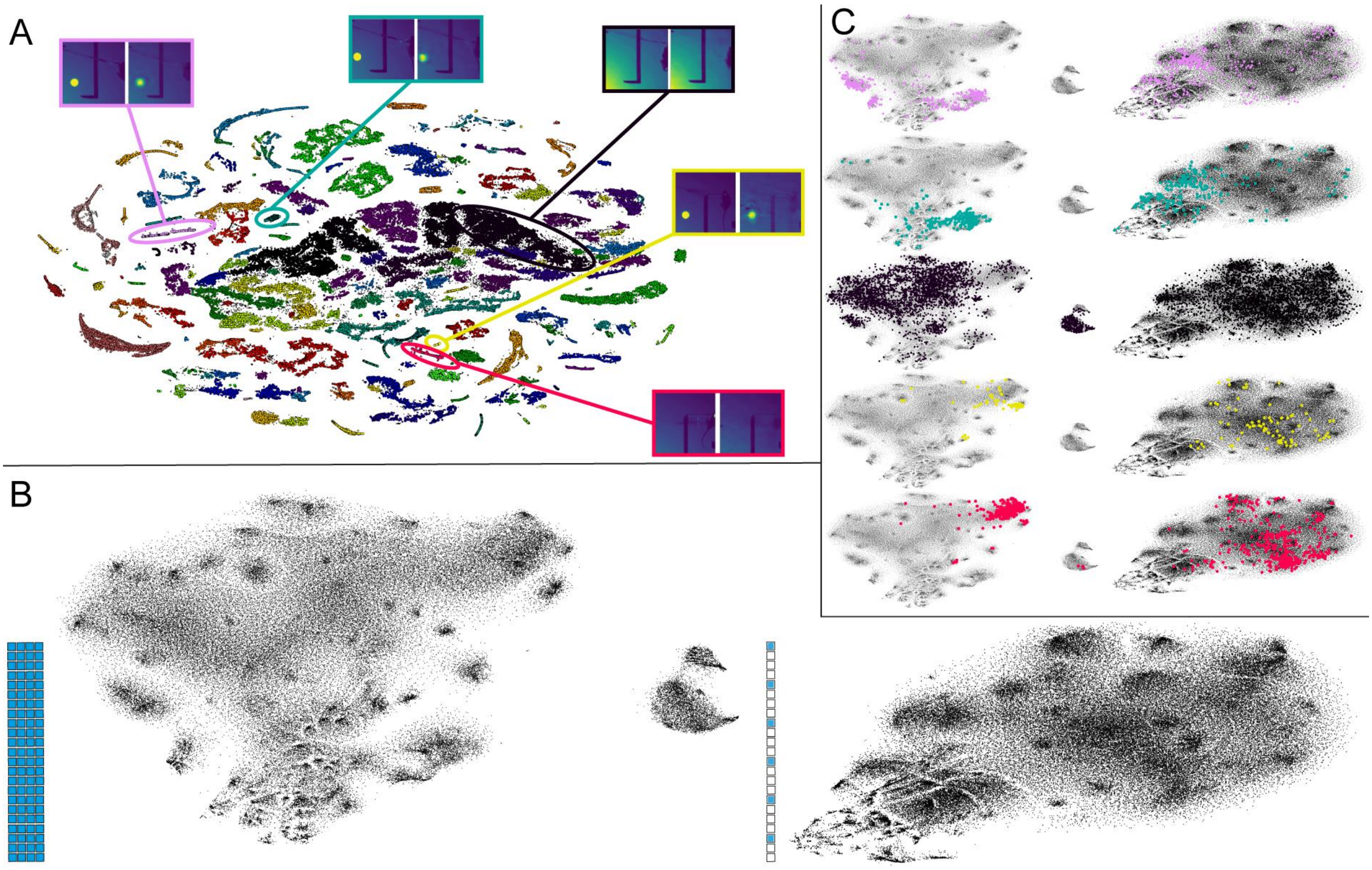


Figure 3. Loss of correspondence between self-similarity of behavior and of the firing rates of recorded neurons with a decrease of the probe's density (see Figure 6) for Animal 3. A, B and C identical to Figure 6.

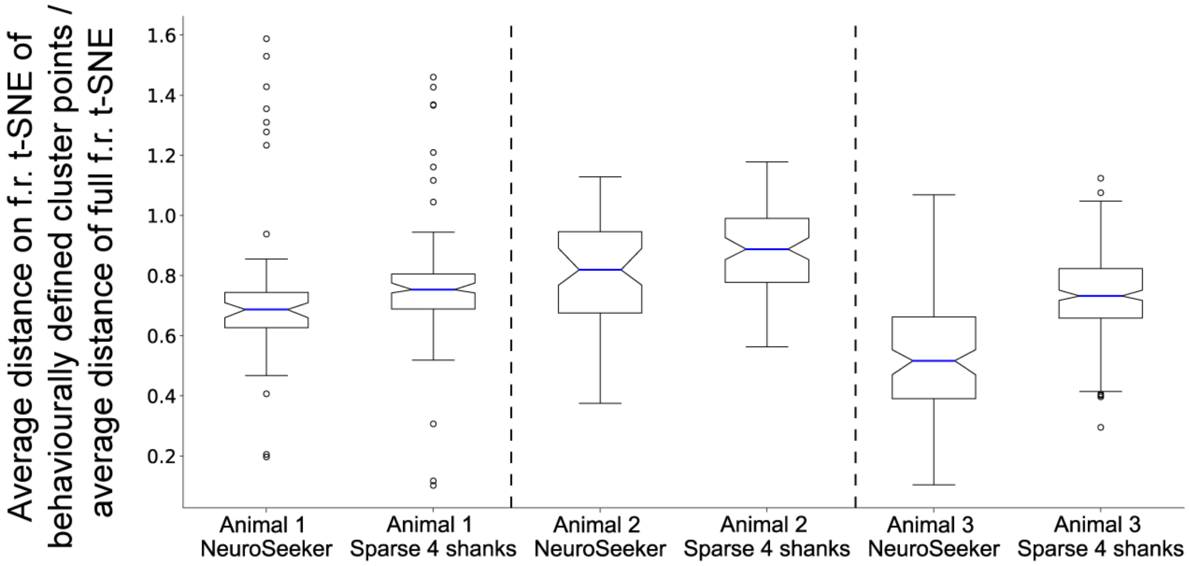


Figure 4. Loss of self-similar patterns over reduction of probe density. Each bar represents a specific recording with either full (NeuroSeeker) or reduced density (Sparse 4 shanks) data set. Each bar shows the statistics generated by a set of points each calculated as the ratio of the average distance of all points on the firing rate t-SNE that correspond to a specific cluster on the behavioral t-SNE over the average distance of 20000 randomly chosen points from the full firing rate t-SNE. So a bar with a lower average (blue line) denotes a recording / probe density set up where the points on the firing rate t-SNE embedding, grouped in the clusters created in the behavioral t-SNE, are more densely packed together.

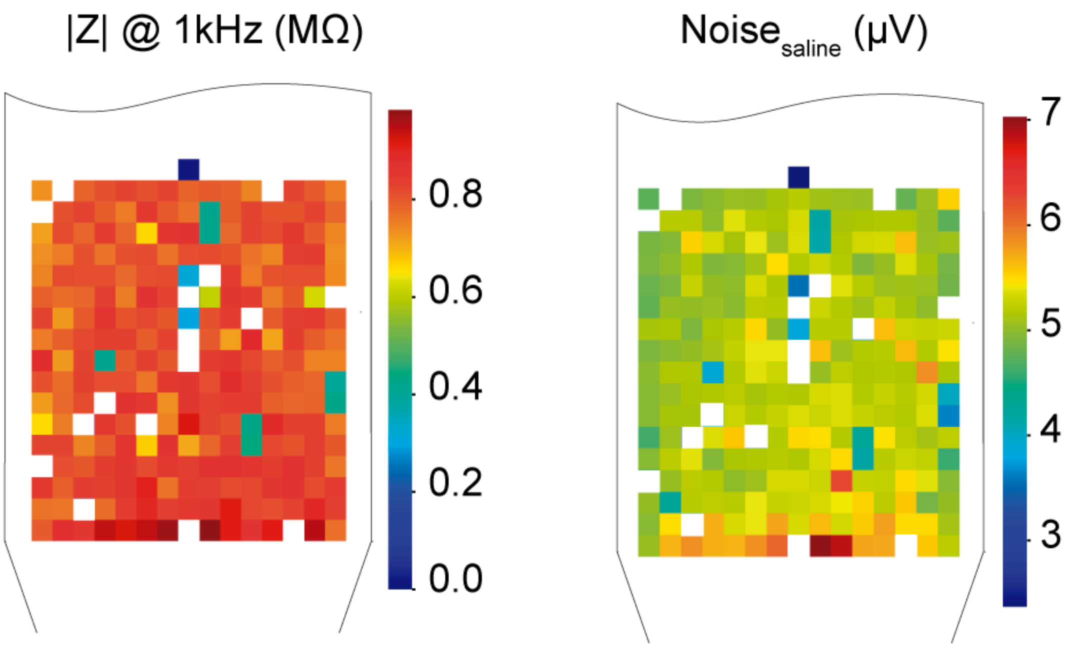


Figure 5. The 256-channel probe: impedance magnitude and noise. Electrical properties of 5 x 5 µm electrodes. Left: the majority of electrodes report an impedance magnitude at 1 kHz lower than 1 MΩ, but 16 of the 255 sites are non-functional because the impedance is higher than 2 MΩ (electrodes represented by white squares). Right: noise magnitude in saline solution. The impedance magnitude and noise measurements were performed with the probe in a dish with saline solution and a reference electrode, Ag-AgCl wire (Science Products GmbH, E-255). The white squares denote the non-functional electrodes.

Table 1. Summary of the dataset used to validate CMOS-based probes. Acute recordings performed with different numbers of active groups, reference type configurations and recording depths. We targeted the brain regions under the stereotaxic coordinates, anterior-posterior -3.4 mm and medial-lateral 1.3 mm. The high-pass cut-off frequency for the AP mode was set to 500 Hz and the low-pass cut-off frequency for the LFP mode was set to 500 Hz. The bias voltage is a parameter that was adjusted for each recording depending mostly on the number of active groups.

| **Filename** | Depth tip (mm) | Number of active groups | Reference type | Time (min) | Bias voltage (V) |
|---|---|---|---|---|---|
| **17_50_36.bin** | 6.7 | 2 | Internal | 1 | 2.25 |
| **17_52_36.bin** | 6.7 | 4 | Internal | 1 | 2.34 |
| **17_54_35.bin** | 6.7 | 6 | Internal | 1 | 2.37 |
| **17_56_25.bin** | 6.7 | 8 | Internal | 1 | 2.39 |
| **17_58_26.bin** | 6.7 | 10 | Internal | 15 | 2.425 |
| **18_15_12.bin** | 6.7 | 12 | Internal | 1 | 2.44 |
| **18_18_41.bin** | 6.7 | 2 | External | 1 | 2.3 |
| **18_20_31.bin** | 6.7 | 4 | External | 1 | 2.38 |
| **18_22_40.bin** | 6.7 | 6 | External | 1 | 2.4 |
| **18_24_20.bin** | 6.7 | 8 | External | 1 | 2.42 |
| **18_26_30.bin** | 6.7 | 10 | External | 15 | 2.43 |
| **18_40_36.bin** | 6.7 | 12 | External | 1 | 2.45 |
| **19_13_16.bin** | 7.6 | 12 | Internal | 15 | 2.43 |

Table 2. Summary of the dataset gathered with the 256-channel probe. Acute recordings (30 minutes long) from anesthetized rats. The recording label specifies the brain region and more specifically the recording position (i.e., anterior-posterior and medial-lateral stereotaxic coordinates and the distance between the brain surface to the tip of the probe).

| **Filename** | Label | Depth tip (mm) | AP (mm) | ML (mm) |
|---|---|---|---|---|
| **amplifier2017-02-08T14_34_33.bin** | Co1 | 0.6 | -3.15 | 1.94 |
| **amplifier2017-02-08T15_34_04.bin** | Co2 | 0.7 | -3.15 | 1.94 |
| **amplifier2017-02-08T16_03_06.bin** | Co3 | 0.8 | -3.15 | 1.94 |
| **amplifier2017-02-08T18_06_19.bin** | H1 | 2.5 | -3.15 | 1.94 |
| **amplifier2017-02-08T18_38_09.bin** | H2 | 3.3 | -3.15 | 1.94 |
| **amplifier2017-02-08T20_04_54.bin** | H3 | 3.5 | -3.15 | 1.94 |
| **amplifier2017-02-08T20_54_26.bin** | T1 | 4.6 | -3.15 | 1.94 |
| **amplifier2017-02-08T21_38_55.bin** | T2 | 6.4 | -3.15 | 1.94 |
| **amplifier2017-02-16T15_37_59.bin** | CR1 | 1.5 | -10.60 | 0.65 |
| **amplifier2017-02-16T16_14_15.bin** | CR2 | 1.7 | -10.60 | 0.65 |
| **amplifier2017-02-16T16_58_01.bin** | CR3 | 2.1 | -10.60 | 0.65 |
| **amplifier2017-02-23T14_38_33.bin** | Co4 | 1.4 | +1.91 | 1.79 |
| **amplifier2017-02-23T15_48_36.bin** | St1 | 3.8 | +1.91 | 1.79 |
| **amplifier2017-02-23T17_29_48.bin** | St2 | 5.5 | +1.91 | 1.79 |
| **amplifier2017-02-23T16_55_00.bin** | St3 | 5.6 | +1.91 | 1.79 |
| **amplifier2017-02-23T18_25_19.bin** | CoP1 | 2.8 | +2.02 | 4.11 |
| **amplifier2017-02-23T19_00_56.bin** | CoP2 | 3.2 | +2.02 | 4.11 |
| **amplifier2017-02-23T19_36_39.bin** | CoP3 | 3.6 | +2.02 | 4.11 |

Table 3. Summary characteristics of the CMOS and the ultra-dense passive probes

| Probe | Electrode type | Amplification | # electrodes | Size of recording area (µm x µm) | Probe thickness (µm) | Electrode size (µm) | Electrode pitch (µm) |
|---|---|---|---|---|---|---|---|
| CMOS | Active, scanned | On-board | 1344 | 8000 x 100 | 50 | 20 x 20 | 22.5 |
| Ultra-dense | Passive | Intan RHD2000 | 255 | 100 x 100 | 50 | 5 x 5 | 6 |
